# Induction of hysteretic chromatin states by CRISPRa reveals locus-specific encoding of transcriptional memory

**DOI:** 10.64898/2026.09.29.755468

**Authors:** Guy C. Bedford, Sangrok Han, Derek L. Yang, Devika Dua, Hakeem Shitta-Bey, Michael A. Beauregard, Andrew R. DiNardo, Isaac B. Hilton, Michael R. Diehl

**Author notes:** Corresponding authors. (I.B.H.); (M.R.D.).

## Abstract

Transcriptional memory is a phenomenon that allows cells to ‘remember’ prior stimulatory history, which can alter subsequent gene expression patterns and shape development, immunity, and disease. Although various epigenetic adaptations have been implicated, it remains unclear whether the mechanics of transcription alone are sufficient to install durable memory independently of complex activation and global phenotypic changes induced by inflammatory and environmental stimuli. Here, we employ CRISPR activation (CRISPRa) to uncouple gene activation from global signaling and test whether memory can be established selectively by a local transcriptional pulse in primary mesenchymal stromal cells (MSCs). Key stimulus-responsive genes (*IDO1, CXCL11*) remain sensitized for weeks after either IFNγ exposure or locus-specific CRISPRa perturbations. Histone PTM profiling links this memory to epigenetic hysteresis involving persistent loss of promoter H3K27me3. Single-molecule chromatin mapping by DAF-seq showed that this adaptation does not coincide with gains in promoter accessibility, but suggests it supports increased transcription factor reacquisition in rechallenged cells. Finally, while H3K27me3 loss appears to be a key hallmark of strongly induced memory genes, multi-omic analyses showed that persistent H3K27me3 hysteresis also occurs at a small subset of non-memory genes. Coordination of H3K27me3 hysteresis with additional cis factors therefore likely shapes transcriptional memory in a locus-specific fashion, underscoring the utility of targeted CRISPRa perturbations for dissecting these responses.

## INTRODUCTION

Activation of intracellular signaling pathways often triggers extensive chromatin remodeling at stimulus-responsive gene loci. These multifaceted biochemical and structural adaptations can persist across successive cell divisions long after the initiating signaling event has resolved (*1*, *2*). The epigenetic “bookmarks” that remain at a gene locus can produce diverse hysteretic transcriptional behaviors, or transcriptional memory, depending on their interactions with other local regulatory factors (*3*), the nature of the stimulus (*4*), and the exposure pattern (*5*). Among the most well characterized memory behaviors are ‘classical’ transcriptional memory (*6–8*), which is marked by accelerated and/or potentiated transcription upon reactivation, and refractory/tolerized behavior (*4*), in which secondary transcriptional outputs are attenuated. Importantly, the epigenetic adaptations underlying these hysteretic responses are often mitotically stable and can endure across multiple rounds of cell division (*9*).

Transcriptional memory occurs across diverse cell types and biological contexts, and is increasingly implicated in human health and disease. For instance, stimulus-responsive epigenetic reprogramming allows innate immune cells to mount faster and more potent responses to repeated inflammatory challenges and/or infections (*10*, *11*). This remodeling has been implicated in mechanisms that confer broad protection against pathogens unrelated to the initial challenge, a phenomenon termed trained immunity. Beyond immune cells, transcriptional memory is associated with the accelerated expression of inflammation-responsive wound-healing genes in epidermal stem cells (*12*, *13*). Memory has also been reported in hematopoietic (*14*), stromal (*7*), and transformed cancer cells (*6*). Given the growing evidence linking memory to chronic inflammation (*15*), autoimmunity (*16*), obesity (*17*), and hyperactive signaling in various other settings, there is a pressing need to stratify the molecular drivers of transcriptional hysteresis at individual loci, both to clarify their roles in cell physiology and to harness memory responses for biomedical applications.

The molecular mechanisms that establish and maintain transcriptional memory are complex and multifaceted. Local cis- and trans-acting factors, as well as broader spatial and global epigenetic changes have both been implicated, including retention of transcription factors at the regulatory elements of memory genes (*12*, *13*), histone post-translational modifications (PTMs) (*18*), altered DNA methylation (*5*, *9*), chromatin topological reorganization (*3*, *6*, *19*), and metabolic rewiring (*20*). Yet, despite the involvement of each of these regulatory layers, and potential reinforcement from feedback among them, the foundational triggers for memory may originate from the mechanics of gene expression itself. Transcription alone is known to alter histone PTMs and remodel local chromatin architecture (*21–23*). Nevertheless, it remains unclear whether primary, transcription-coupled events are sufficient to install memory independently, or whether durable memory is also contingent upon the activation of downstream responses of composite signaling pathways (*24*, *25*).

Here, we leverage transient mRNA delivery of CRISPR activation (CRISPRa) (*26*) complexes to facilitate precision-targeted expression perturbations and demonstrate that local activation of transcription is sufficient to install persistent chromatin adaptations and generate transcriptional memory. Using the responses of primary human mesenchymal stromal cells (MSCs) to sequential exposures of the inflammatory cytokine interferon-γ (IFNγ) as an experimental model, we first show that multiple functionally significant interferon stimulated genes (ISGs) exhibit both accelerated and potentiated expression upon secondary IFNγ exposure. Key examples include *IDO1*, which encodes indoleamine 2,3-dioxygenase 1, a potent immunomodulatory effector and important marker of MSC therapeutic potency (*27*, *28*), and *CXCL11*, which encodes the C-X-C motif chemokine ligand 11. Both genes remained sensitized towards IFNγ rechallenge for at least 21 days (∼5 cell divisions). Similar classical memory behavior occurs selectively at both gene loci using mRNA delivery of CRISPRa tools as a locally-targeted primary activation signal, but importantly, without the multifaceted alterations and transcriptional drift that occur upon activation of the IFNγ pathway. Cleavage Under Targets and Release Using Nuclease (CUT&RUN) profiling of histone PTMs suggests that memory at these highly responsive ISGs is linked to a persistent and progressive depletion of repressive H3K27me3 marks at their promoters. This depletion was notably observed when either IFNγ or CRISPRa was used as the primary stimulant, indicating that local transcription alone, in the absence of global IFNγ-induced perturbations, is sufficient to establish this hysteretic chromatin signature. Nevertheless, a small subfraction of non-memory ISGs also exhibits hysteretic reduction in H3K27me3 without appreciable transcriptional hysteresis, indicating that potential coupling of H3K27me3 to other regulatory features are necessary to fully imprint transcriptional memory.

## RESULTS

### *IDO1* displays robust transcriptional memory in response to IFNγ stimulation

We chose to study memory in adipose-derived, primary human MSCs because these cells possess normal karyotypes, exhibit epigenetic plasticity, and have high sensitivity towards IFNγ and other proinflammatory signals. Furthermore, inflammatory transcriptional memory has not been rigorously demonstrated in these cells. We investigated memory by first quantifying IDO1 protein expression in response to sequential primary and secondary stimulation with IFNγ. Early-passage (2) MSCs were thawed, expanded once, treated with IFNγ for 24 hrs, and then subcultured for at least 10 days, over which cells underwent approximately 2.3 cumulative population doublings (2.3 ± 0.1 for IFNγ-treated versus 2.8 ± 0.1 for untreated cells; mean ± SEM; Fig. S1A), prior to secondary stimulation. Expression behaviors were compared across four conditions: unstimulated cells (Naive), cells exposed to an initial stimulation (Stimulated), cells that were stimulated and then rested for a ten-day period (Rested), and cells re-stimulated a second time after the rest period (Rechallenged) (**Fig. 1A**). Intracellular IDO1 protein and mRNA levels were quantified across conditions using flow cytometry, western blotting and RNA sequencing. While exhibiting high activation levels (>450-fold) upon IFNγ exposure, IDO1 expression returned to undetectable levels following a 10-day rest period (Fig. 1B). Memory was reflected in comparison of IDO1 expression in stimulated and rechallenged cells. Here, rechallenged MSCs displayed both a 40% increase in the number of IDO1-positive cells after IFNγ exposure (70.2 ± 0.9% versus 98.5 ± 0.1%), as well as a 1.65-fold increase in mean fluorescence intensity (MFI) within the IDO1-positive population (4,834 ± 64 versus 7,961 ± 126; **Fig. 1B, Fig. S1B**).

**Fig. 1.**
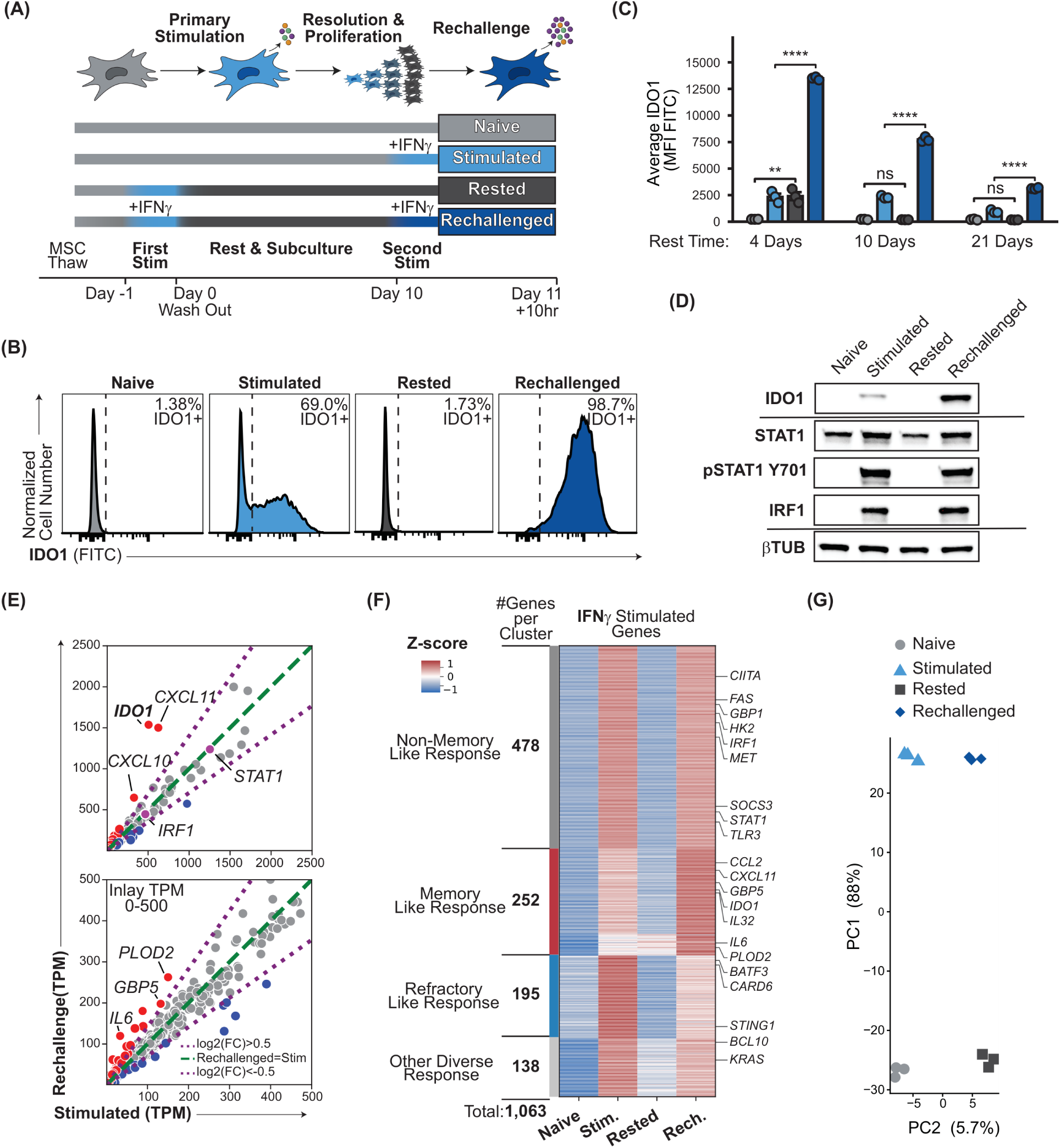
IFNγ induces diverse transcriptional memory trajectories in MSCs. **(A)** Schematic of the inflammatory-memory paradigm in MSCs, with the experimental timeline and the four conditions compared throughout: Naive (no IFNγ), Stimulated (IFNγ on day 10 only), Rested (IFNγ on day 0, then rest), and Rechallenged (IFNγ on day 0 and again on day 10). **(B)** Flow-cytometry histograms of intracellular IDO1 across conditions 10hr post rechallenge; dashed line, IDO1+ gate. **(C)** Mean intracellular IDO1 protein by flow cytometry (MFI) across conditions at three rest durations before rechallenge (4, 10, and 21 days). **(D)** Western blots across conditions for IDO1, total STAT1, phospho-STAT1 (Y701), and IRF1; β-tubulin (βTUB), loading control. **(E)** Per-gene correlation of Rechallenged versus Stimulated expression (TPM); inset magnifies the 0–500 TPM range. Green dashed line, Rechallenged = Stimulated; purple dotted lines, log2(fold change) = ±0.5. **(F)** K-means heat map (k = 5) of z-scored expression for 1,063 IFNγ-stimulated genes (rows) across conditions (columns), with the five clusters grouped into four response classes (Non-Memory-Like, Memory-Like, Refractory-Like, Other Diverse) with representative genes annotated. **(G)** Principal-component analysis of the RNA-seq data (PC1, 88%; PC2, 5.7%). In C, bars are mean ± SEM with individual replicates (n = 3 biological replicates); significance by one-way ANOVA with Tukey’s post-test. **** P < 0.0001, *** P < 0.001, ** P < 0.01, * P < 0.05; ns, not significant.

*IDO1* memory dynamics were further evaluated by altering IFNγ exposure (**Fig. S1C**) and varying the total time of the rest period (**Fig. 1C**). Comparisons across IFNγ exposure times demonstrated *IDO1* activation occurs both more rapidly and reaches higher saturation levels after rechallenge compared to primary stimulation. This accelerated and potentiated reactivation was independently validated in two other donor lines by western blotting (**Fig. S1D**). Moreover, analyses across different rest period durations revealed memory persists for up to 21 days, although the overall magnitude of IDO1 induction decreased with increasing rest time (**Fig. 1C**). Consistent with observations reported in other cell systems (*24*), the core activating transcription factors in the IFNγ pathway (STAT1 and IRF1) do not display memory (**Fig. 1D**), indicating that these memory effects are not attributable to sustained STAT1 or IRF1 expression after initial IFNγ exposure.

### IFNγ induces diverse transcriptional memory trajectories with some transcriptional drift

We next performed a series of RNA sequencing analyses to identify the full suite of stimulus-responsive genes that display transcriptional memory in MSCs. Primary IFNγ stimulation was found to upregulate a suite of 1,063 protein-coding interferon stimulated genes (ISGs), the composition of which matched well with canonical activation profiles within the IFNγ pathway (*29*). Although this broad activation response largely returned to the baseline of naive cells following a ten-day rest period (gene set enrichment for “response to type II interferon”: stimulated versus naive NES = 2.36, p = 1.8 × 10⁻⁸; rested versus naive NES = 1.28, p = 0.26) (**Fig. S2A**), many ISGs exhibited some form of transcriptional hysteresis upon rechallenge (266 of 1,063 genes differentially expressed between the stimulated and rechallenged conditions, DESeq2 FDR < 0.05). *IDO1* was among a collection of ISGs that exhibited a highly pronounced memory effect; followed closely by *CXCL11* (**Fig. 1E**). Both ISGs exhibited high levels of transcriptional activation in response to primary stimulation (Δ Transcripts Per Million (TPM) <2 to >500; 453-fold and 521-fold for *IDO1* and *CXCL11*, respectively), and substantially potentiated activation after rechallenge, reaching >1,500 TPM, a further 3.0- and 2.4-fold above primary stimulation. An additional set of 48 ISGs were found to display ‘classical’ memory behavior, marked by a log2 fold change > 0.5 between the stimulated and rechallenged conditions. The identification of nearly 50 distinct classical memory genes indicates this type of hysteretic behavior occurs in a sizable subset of IFNγ-responsive genes (**Fig. S2B**).

Consistent with previous studies (*7*), unsupervised K-means clustering using z-score– normalized expression values for all 1,063 ISGs further revealed ISGs can exhibit a spectrum of different dynamic behaviors (**Fig. 1F**). The largest cluster contained 478 genes that did not display hysteretic behavior. Two additional clusters (n= 201 and 51) exhibited memory behaviors in which expression was highest in the rechallenged condition. Refractory behavior as well as more complicated hysteretic responses that do not fit the commonly described definitions of memory were also observed (n = 195 and 138). The diversity of transcriptional hysteresis was not unique to IFNγ stimulation, as primary stimulation and rechallenging with an alternative proinflammatory cytokine, TNFα, also yielded a spectrum of similar hysteretic responses, although refractory response appeared to be favored over ‘classical memory’ in this case (n = 472 and 158 genes, respectively) (**Fig. S3A-C**).

We next examined transcriptional drift in the composite, the genome-wide transcriptional profile of MSCs across experimental conditions due to IFNγ stimulation. While MSCs retain their identity across all experimental conditions with respect to expression marker panels defined by the International Society for Cellular Therapy (**Fig. S4A**), a substantial number of genes were differentially expressed between the naive and rested conditions (1030 genes; 548 up, 482 down, |log2FC| > 0.5, adjusted P < 0.05) (**Fig. S4B**). Due to the potency of IFNγ activation, principal component analysis (PCA) of the MSC transcriptome showed that stimulation was by far the dominant source of transcriptional variation, resulting in pronounced separation between either stimulated conditions (stimulated and rechallenged) and unstimulated conditions (naive and rested) (**Fig. 1G**). Although a smaller change, naive and rested cells were nevertheless separated along PC2 (5.7% of variance). Stimulated and rechallenged cells showed similar separation along the PC2 axis, indicating this transcriptional drift is persistent (**Fig. 1G**). Gene set enrichment of naive vs rested cells returned 293 GO biological processes at FDR < 0.05, with extracellular matrix organization among those most enriched in rested cells (NES = 2.13, FDR = 1.5 × 10⁻⁷) and cytoplasmic translation among those most depleted (NES = −2.15, FDR = 5.3 × 10⁻⁶) (**Fig. S4C**). Similar net transcriptional drift was observed between the naive and rested cells after TNFα stimulation, although the variability across groups appears to be smaller than the drift observed with IFNγ (364 genes; 178 up, 186 down; **Fig. S3C**).

In sum, these analyses show that ISGs display diverse types of temporal hysteretic behaviors in response to repeated IFNγ inflammatory cytokine challenge. Moreover, even though ISG activation largely resolves to near baseline during extended rest periods, residual transcriptional drift occurs between naive and rested cells, which likely arises from the highly complex and multifaceted transcriptomic, proteomic and metabolic alterations that contribute to inflammatory cytokine signaling pathways.

### mRNA delivery of CRISPRa tools introduces targeted, transient expression perturbations

To circumvent the challenges stemming from the complexity of biochemical signaling pathway activation, resultant diversity of memory trajectories and transcriptomic drift, we chose to employ CRISPRa mediated transcriptional activation as a route to selectively drive expression at defined loci, in the absence of cytokine exposure. CRISPRa targeting was achieved via cotransfection of chemically synthesized single guide RNAs (sgRNAs) with *in vitro* transcribed (IVT) mRNA encoding for the CRISPRa effector VP64-dCas9-VP64. Importantly, VP64 acts through recruitment of endogenous coactivators rather than direct catalytic deposition of histone modifications (*30–32*), isolating transcription as the perturbation.

Analyses with a control dCas9-eGFP construct demonstrated that MSCs can be transfected more efficiently using mRNA delivery (>89% GFP positive cells) compared to DNA plasmid delivery (1% GFP positive cells) (**Fig. S5A**). Efficient transcriptional activation (∼90% of cells) was also achieved in MSCs using a VP64-dCas9-VP64 construct with an sgRNA targeting *CD34*, which served as a CRISPRa benchmark standard that is not detectably expressed in native adipose-derived MSCs (**Fig. S5B**). Targeting VP64-dCas9-VP64 to a single site in the *IDO1* promoter yielded a modest but reproducible increase in intracellular IDO1 protein at 72 hours post-transfection (geometric MFI increased >1.6-fold over a non-targeting). VP64-dCas9-VP64 yielded lower activation levels of IDO1 compared to IFNγ (**Fig. S5D,E**). Nevertheless, this activation occurred without driving STAT1 Y701 phosphorylation, confirming the approach bypasses canonical IFNγ signaling (**Fig. S5C**).

Importantly, IDO1 activation was transient and decayed to negligible levels in rested cells, demonstrating that mRNA delivery of VP64-dCas9-VP64 to the *IDO1* loci facilitates selectively targeted, transient transcriptional perturbations for dissecting memory behaviors at a defined locus (**Fig. 2A,B**). Temporal western blot analysis of VP64-dCas9-VP64 and IDO1 protein expression display detectable VP64-dCas9-VP64 signals within 4 hours of mRNA delivery transfection, and a strong decay to an undetectable level by day 3. IDO1 expression levels lagged behind VP64-dCas9-VP64 and peaked between days 4 to 5. This paralleled the biochemical IFNγ stimulation of IDO1 (**Fig. 2C,D**).

**Fig. 2.**
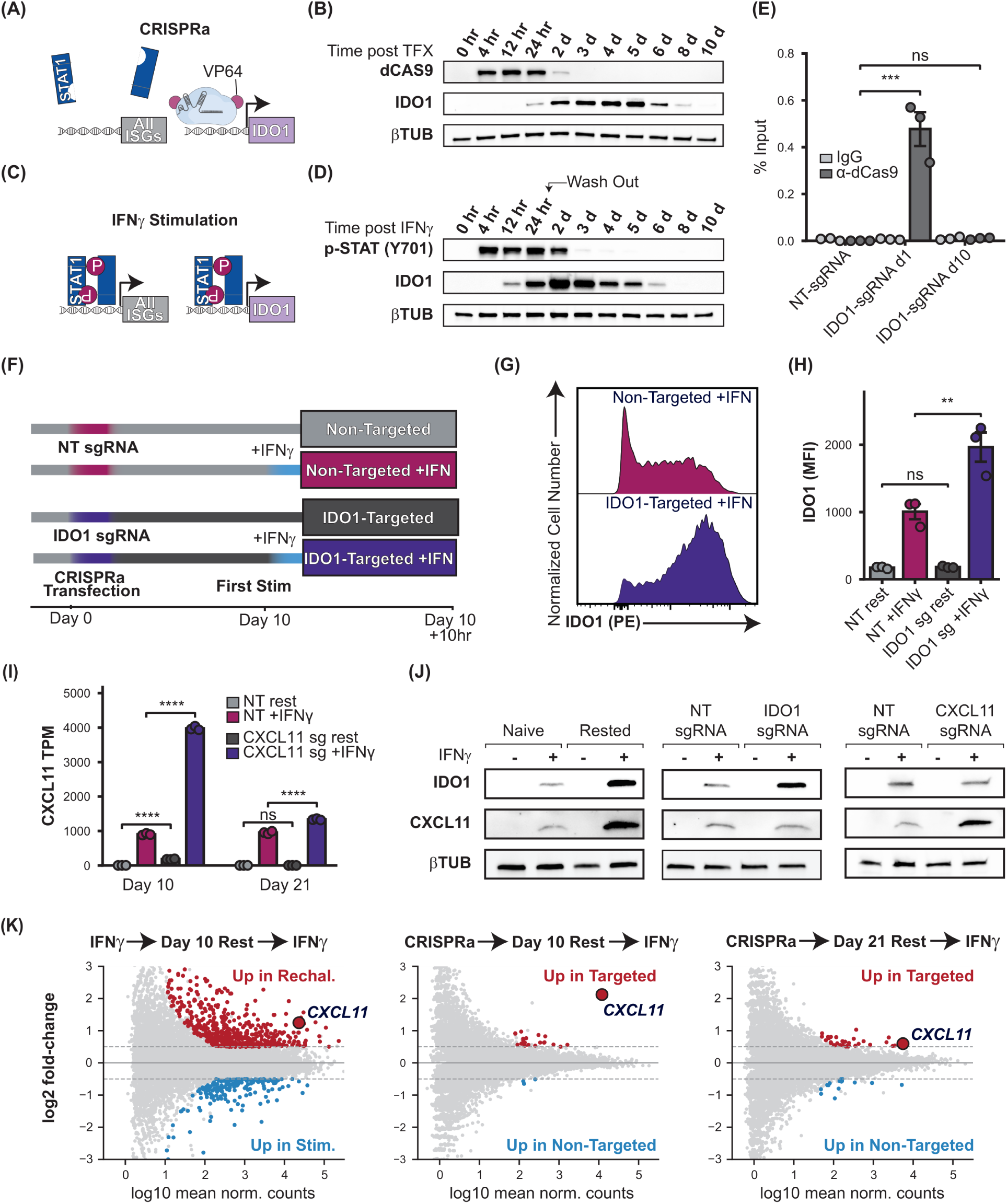
Transient mRNA CRISPRa installs locus-specific synthetic transcriptional memory at *IDO1* and *CXCL11* without genome-wide perturbation: **(A)** Schematic of mRNA CRISPRa: a VP64–dCas9–VP64 effector targeted to the *IDO1* promoter drives *IDO1* selectively. **(B)** Western blot time course after transfection of MSCs with VP64–dCas9–VP64 mRNA plus *IDO1*-targeting sgRNA (0 hr–10 d), probed for dCas9, IDO1, and βTUB. **(C)** Schematic of canonical IFNγ stimulation. **(D)** Western blot time course after IFNγ stimulation (0 hr–10 d; arrow, cytokine wash-out at 24 hr), probed for phospho-STAT1 (Y701), IDO1, and βTUB. **(E)** ChIP-qPCR of VP64-dCas9-VP64 occupancy at the *IDO1* promoter (% input) for NT-sgRNA, IDO1-sgRNA day 1, and IDO1-sgRNA day 10; IgG control versus α-dCas9. (F) Experimental timeline for CRISPRa priming: MSCs transfected on day 0 with a non-targeting (NT) or *IDO1*-targeting sgRNA, rested to day 10, then given a first IFNγ stimulation and harvested at day 10 + 10 hr. Four conditions: Non-Targeted, Non-Targeted +IFNγ, IDO1-Targeted, IDO1-Targeted +IFNγ. (G) Flow-cytometry histograms of intracellular IDO1 after the first IFNγ stimulation for Non-Targeted +IFNγ (top) and IDO1-Targeted +IFNγ (bottom). **(H)** IDO1 mean fluorescence intensity (MFI) by flow cytometry across the four conditions in F. (I) *CXCL11* transcript abundance (TPM, RNA-seq) at two rest durations (Day 10, Day 21) for NT rest, NT +IFNγ, CXCL11 sg rest, and CXCL11 sg +IFNγ. **(J)** Western blots for IDO1, CXCL11, and βTUB (loading) ± IFNγ. Left, IFNγ training conditions; middle, *IDO1* CRISPRa priming; right, *CXCL11* CRISPRa priming. **(K)** MA plots of RNA-seq differential expression (log2 fold change versus mean normalized counts); significant DEGs (FDR < 0.05, |log2FC| > 0.5) colored by direction of change (red, up; blue, down), CXCL11 labeled, dashed guides at log2FC ±0.5. Left, IFNγ training (Rechallenged versus Stimulated, Day 10); middle, CRISPRa priming (CXCL11-Targeted +IFNγ versus Non-Targeted +IFNγ, Day 10); right, the same contrast at Day 21. Bars, mean ± SEM with individual replicates (n = 3 biological replicates). Significance: E, one-way ANOVA with Dunnett’s post-test versus NT-sgRNA; H, two-way ANOVA (guide × IFNγ); I, gene-level DESeq2 adjusted P (Wald test, Benjamini–Hochberg FDR). **** P < 0.0001, *** P < 0.001, ** P < 0.01, * P < 0.05; ns, not significant.

Finally, to further confirm that VP64-dCas9-VP64 no longer occupied the *IDO1* promoter, we performed chromatin immunoprecipitation qPCR (ChIP-qPCR) for dCas9 at days 1 and 10 post-transfection. At day 1 post-transfection, ChIP-qPCR demonstrated robust VP64-dCas9-VP64 promoter occupancy relative to input. By day 10, however, promoter binding was no longer detectable (**Fig. 2E**). Together these data demonstrate that mRNA delivery of VP64-dCas9-VP64 in MSCs can enable selective activation and resolution of ISGs for probing memory mechanisms.

### CRISPRa installs locus-specific transcriptional memory without genome-wide perturbation

We next tested whether transient and locus-specific activation could sensitize the *IDO1* locus to subsequent IFNγ challenges (**Fig. S5F**). Here, MSCs were transfected with either an sgRNA targeting *IDO1* or a non-targeting control sgRNA and allowed to rest for ten days until *IDO1* expression had returned to baseline. Both CRISPR activated groups were then rechallenged with IFNγ stimulation (**Fig. 2F**). Cells in which *IDO1* had previously been activated via CRISPRa exhibited significantly more potent IDO1 induction upon IFNγ stimulation, reaching roughly twice the IDO1 protein expression level of cells that received a non-targeting sgRNA (MFI = 1007 ± 113 vs MFI = 1966 ± 218; mean ± SEM, for non-targeted+ IFNγ and *IDO1*-targeted+ IFNγ respectively; **Fig. 2G,H**). Overall, the accelerated rate and enhanced magnitude of this response mirrored the accelerated and potentiated kinetics of transcriptional memory observed with IFNγ alone, indicating that transient, locus-specific CRISPRa is sufficient to establish a similar sensitized state at the *IDO1* locus (**Fig. 2G,H and Fig. 1B**).

The ability to install transcriptional memory via CRISPRa was not unique to *IDO1*. The ISG *CXCL11*, which also displays robust transcriptional memory to multiple challenges with IFNγ (**Fig. 1E, Fig. S5G**), was also sensitized when the locus was targeted with CRISPRa (**Fig. 2I, Fig. S5H**). Ten days after mRNA transfection with a *CXCL11*-targeting sgRNA, the *CXCL11* response to IFNγ was potentiated 4.4-fold relative to the non-targeting control (3979 ± 33 versus 904 ± 15 TPM, mean ± SEM). With a longer rest of 21 days the memory effect fell to 1.4-fold but still produced a significant enhancement over the non-targeting control (1336 ± 13 versus 955 ± 21 TPM). Interestingly, at day 10, *CXCL11* transcripts also remained elevated in rested cells in the absence of any IFNγ stimulation (182 ± 5 versus 1.6 ± 0.4 TPM). This residual expression resolved completely by day 21 (1.6 ± 0.4 versus 1.5 ± 0.4 TPM), indicating that the enhanced rechallenge response at that timepoint reflects a primed but transcriptionally silent locus rather than carryover of the initial activation.

In contrast to biochemical stimulation, activation by VP64-dCas9-VP64 was highly selective and localized to the specific sgRNA target locus. For instance, western blot analysis confirmed targeting was locus-specific with no activation crosstalk between *IDO1* and *CXCL11* (Fig. **2J**). In addition, while *IDO1* targeting was found to perturb MSC viability, reducing viability to 82.8 ± 3.2% of the non-targeting control at 72 hr (one-way ANOVA with Dunnett’s test, p = 0.042, n = 3), targeting *CXCL11* had no effect on viability (100.4 ± 4.3% of control, p = 0.996) (**Fig. S6B**). IDO1 catabolizes tryptophan, and converts intracellular and medium tryptophan into kynurenine, with both positive and negative feedback loops (*33*). In contrast, CXCL11 is a secreted chemokine acting on CXCR3, which MSCs do not express (**Fig. S6C**), which should reduce if not eliminate downstream signaling feedback.

Bulk RNA-sequencing further confirmed that this reciprocal specificity extends to the transcriptome, demonstrating that CRISPRa-induced memory at *CXCL11* is decoupled from stress responses and genome-wide transcriptional changes that accompany IFNγ priming. For example, few differentially expressed genes (DEGs) were identified between *CXCL11*-targeted and non-targeted cells upon IFNγ rechallenge (25 DEGs at day 10 and 51 DEGs at day 21, with *CXCL11* among the most significant) (**Fig. 2K**). Similarly small background perturbations were identified when comparing rested states (27 DEGs at day 10 and 53 DEGs at day 21) (**Fig. S6A**). Together, these data indicate CRISPRa can install synthetic transcriptional memory through a locus-specific mechanism wherein memory-encoding information is strictly constrained in *cis* to the target locus, while producing minimal global perturbations or off-target effects.

### H3K27me3 dynamics instruct transcriptional memory at a subset of memory genes

Histone PTMs at stimulus-responsive loci change substantially upon activation with IFNγ and many other signaling pathways and have been implicated as drivers of memory (*7*, *18*, *34*). We therefore probed whether signatures of transcriptional memory are encoded by alterations of histone PTMs at the promoters of stimulus-responsive genes in MSCs. A suite of 4 PTM marks was characterized through a combination of CUT&RUN qPCR and sequencing. These marks included H3K4me3, H3K36me3, and H3K36me2, which mark active promoters, transcribed gene bodies and intergenic regions respectively (*35*, *36*), and H3K27me3, which marks facultative heterochromatin (*37*). We chose these marks due to their previous association with transcriptional memory in other cells (*7*, *18*, *25*) as well as their strong coupling to the transcriptional activity of RNA Polymerase II (RNAPII) across distinct phases of the transcription cycle (*21*). Specifically, H3K4me3 serves as a canonical marker of active or primed promoters, whereas H3K36me2 and H3K36me3 track actively elongating Pol II across gene bodies and are known to directly antagonize the propagation of polycomb-mediated repression. By assessing the dynamics of these marks, we sought to determine whether local transcriptional elongation results in persistent epigenetic bookmarks capable of sustaining locus-specific memory during extended rest periods.

Histone PTMs were mapped across the full training paradigm (naive, IFNγ-stimulated, rested, and IFNγ-rechallenged) for H3K27me3 and H3K4me3 and across naive, IFNγ-stimulated for the H3K36me2/3 marks. Clear patterns and memory effects were observed across these PTMs, particularly at *CXCL11* and *IDO1*. The CUT&RUN tracks for *CXCL11* and *IDO1* are displayed in **Figs. 3A,B** and **S7A,B**. H3K4me3 (top right), which is known to be tightly coupled to promoter accessibility and transcription initiation, was elevated at the gene promoters of both loci in IFNγ stimulated conditions, and further potentiated in rechallenged cells, consistent with the transcriptional memory effect observed via RNAseq. However, H3K4me3 signals remain at baseline levels in both naive and rested cells. H3K36me3 (bottom left) and H3K36me2 (bottom right), marks associated with active RNA polymerase II elongation showed strong induction from naive to stimulated.

**Fig. 3.**
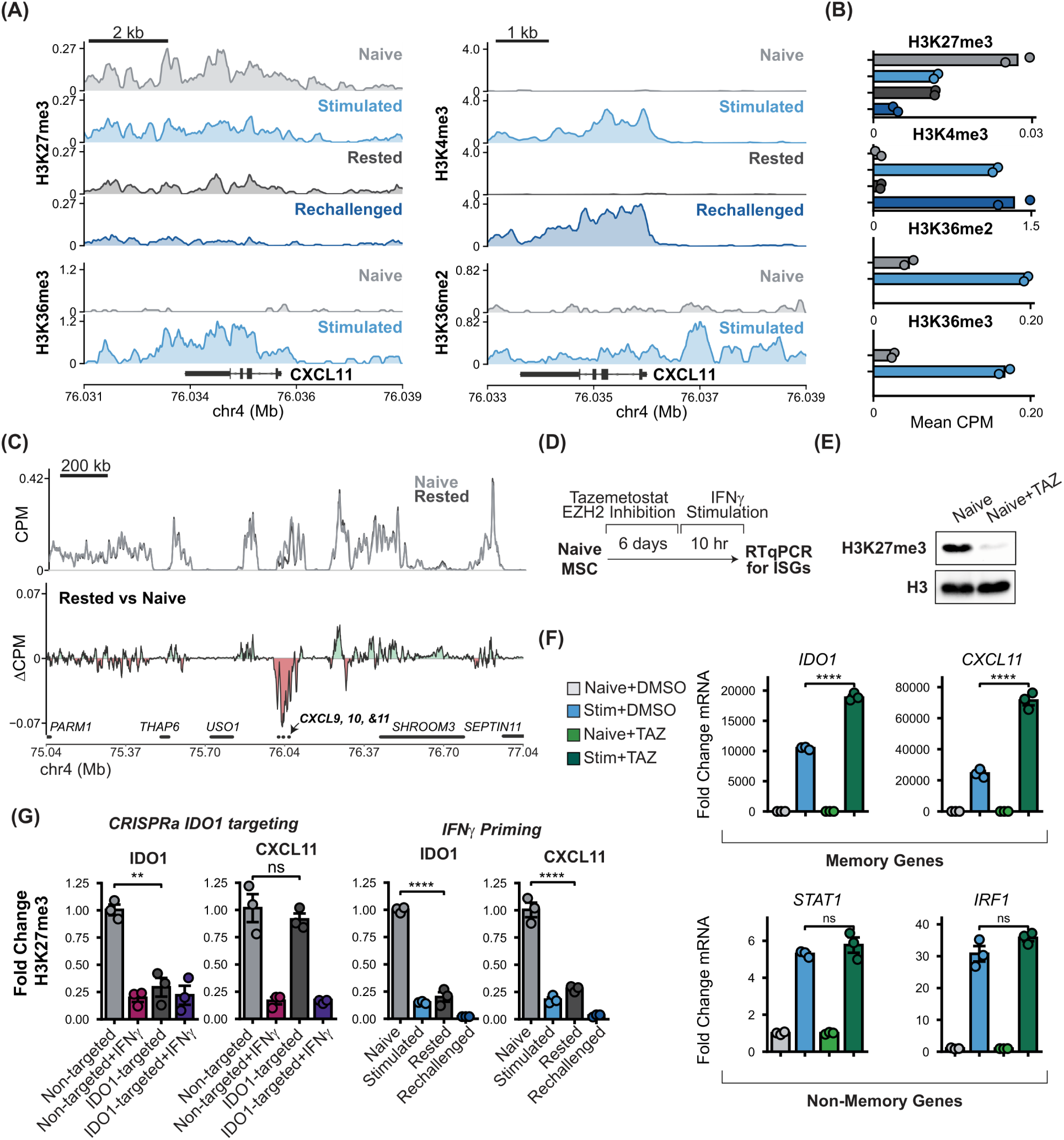
H3K27me3 dynamics instruct transcriptional memory at *IDO1* and *CXCL11*: **(A)** Genome-browser CUT&RUN tracks at the *CXCL11* locus. Left, H3K27me3 (Naive, Stimulated, Rested, Rechallenged) and H3K36me3 (Naive, Stimulated). Right, H3K4me3 (four conditions) and H3K36me2 (Naive, Stimulated). **(B)** Mean CUT&RUN signal (CPM) at the shown *CXCL1*1 locus for each mark, colored by condition n=2. **(C)** H3K27me3 CUT&RUN across a ±1 Mb window on chr4 (75.04–77.04 Mb) surrounding *CXCL11*. Top, mean coverage overlaying Naive (grey) and Rested (dark grey). Bottom, difference track (Rested – Naive): red, H3K27me3 lost in rested; green, gained in rested. **(D)** Schematic of the EZH2-inhibition experiment: Naive MSCs treated with tazemetostat (TAZ) for 6 days, stimulated with IFNγ for 10 hr, then ISGs measured by RT-qPCR. **(E)** Western blot confirming global H3K27me3 depletion (Naive versus Naive+TAZ); total H3, loading control. **(F)** RT-qPCR fold-change mRNA for memory genes *IDO1*, *CXCL11* and non-memory genes *STAT1*, *IRF1* across Naive+DMSO, Stim+DMSO, Naive+TAZ, and Stim+TAZ. **(G)** CUT&RUN-qPCR of H3K27me3. Left (CRISPRa IDO1 targeting), *IDO1* and *CXCL11* across Non-targeted, Non-targeted +IFNγ, *IDO1*-targeted, and *IDO1*-targeted +IFNγ (normalized to Non-targeted = 1.0). Right (IFNγ priming), *IDO1* and *CXCL11* across Naive, Stimulated, Rested, and Rechallenged (normalized to Naive = 1.0). Bars, mean ± SEM with individual replicates (n = 3 biological replicates). Significance: F, two-way ANOVA (stimulation × tazemetostat); G, two-way ANOVA (guide × IFNγ; CRISPRa targeting) and one-way ANOVA with Tukey’s post-test (IFNγ priming). **** P < 0.0001, *** P < 0.001, ** P < 0.01, * P < 0.05; ns, not significant.

H3K27me3 (top left) exhibited distinct hysteretic dynamics compared to H3K4me3 and H3K36me2/3. Resolving these dynamics required greater sequencing depth than the other marks profiled (**Fig. S8**). In naive cells, H3K27me3 was enriched at the *CXCL11* and *IDO1* promoters relative to non-memory genes such as *STAT1* and *IRF1*, reflecting a repressed baseline state (**Fig. S7C**). Primary IFNγ stimulation reduced H3K27me3 at both loci. Following 10 days of rest, H3K27me3 remained depleted at the *IDO1* and *CXCL11* promoters. Spatial analysis of a 2 Mb region surrounding *CXCL11 (***Fig. 3C**) showed that this rest-associated depletion was confined to its locus, while neighboring polycomb domains appeared to remain intact. The derepressed domain spanned *CXCL11* along with its adjacent, co-regulated memory genes *CXCL9* and *CXCL10*, all of which are potently induced by IFNγ and are among the 50 genes we define as memory genes (**Fig. S7D**).

To determine whether persistent H3K27me3 depletion functions as a molecular driver of memory, we pharmacologically depleted H3K27me3 in naive MSCs using the EZH2 inhibitor tazemetostat prior to IFNγ challenge (**Fig. 3D,E**). Global reduction of H3K27me3 significantly potentiated IFNγ-induced expression of *IDO1* and *CXCL11* (**Fig. 3F**), effectively phenocopying cytokine-induced memory, whereas the responses of non-memory controls, *STAT1* and *IRF1*, remained unaffected (**Fig. 3F**).

Finally, we asked whether selective activation of memory genes via targeted CRISPRa was sufficient to induce a persistent and selective reduction in H3K27me3. MSCs were co-transfected with VP64-dCas9-VP64 and either an *IDO1*-targeting or a non-targeting sgRNA, rested for 10 days as in **Fig. 2F**, then either left untreated or rechallenged with IFNγ. H3K27me3 was measured by CUT&RUN-qPCR at *IDO1* and *CXCL11* (**Fig. 3G**), alongside IFNγ-treated cells that were trained as illustrated in **Fig. 1A**, without CRISPRa. *IDO1*-targeting CRISPRa reduced H3K27me3 at *IDO1* roughly four-fold relative to the non-targeting sgRNA, and yielded comparable levels to cells that were rested after IFNγ stimulation. Furthermore, unlike global IFNγ treatment, targeting *IDO1* via CRISPRa selectively depleted H3K27me3 levels at the *IDO1* locus while leaving H3K27me3 marks at *CXCL11* unaltered (**Fig. 3G**).

### Promoter accessibility tracks stimulation but is not retained between IFNγ exposures

Given the depletion of H3K27me3 at the *IDO1* and *CXCL11* promoters, we used DAF-seq (deaminase-assisted single-molecule fiber sequencing) to test whether this memory effect was associated with increased chromatin accessibility and altered TF localization at these promoters. DAF-seq utilizes enzymatic cytosine deamination on single molecules of DNA to map chromatin accessibility across individual, kilobase-scale DNA fibers at near-single-nucleotide resolution (*38*). The precision gained by these capabilities opens important opportunities to resolve locus-level epigenetic heterogeneity, discriminate distinct subpopulation states, and precisely delineate transcription factor (TF) occupancy patterns.

DAF-seq was performed on targeted genomic regions spanning the promoters of *IDO1* (∼4.1 kb), *CXCL11* (∼3.7 kb), and the non-memory gene *STAT1* (∼2.9 kb). Modification sensitive patches (MSPs) denoting local accessible DNA regions were identified using the DAF-seq Hidden Markov Models (HMM) (**Fig. 4A**). MSP calls at the TSS of *IDO1* and *CXCL11* in naive MSCs were comparable to background levels across both amplicons (4.6% and 12.3% mean MSP-covered area for *IDO1* and *CXCL11*, respectively), consistent with the repressed transcriptional state of these genes in the absence of stimulation. In contrast, the TSS of *STAT1* displayed large accessible domains across ∼97% of fibers. Total MSP density at *STAT1* was also substantially higher (∼33%) compared to *IDO1* (∼15%) or *CXCL11* (∼18%) (**Fig. 4B**). The *IDO1* and *CXCL11* promoters both gained substantial accessibility upon primary IFNγ stimulation. K-means clustering (k = 2) across all fibers for each promoter resolved two clear accessibility states, classified as MSP-rich (open) or MSP-poor (closed) subpopulations (**Fig. 4C**). IFNγ stimulation increased the MSP-rich fraction from 5% to 38% at *IDO1* and from 12% to 85% at *CXCL11*. Fiber accessibility reverted nearly completely back to the naive condition in rested MSCs, with only small residuals between rested and naive conditions (*IDO1* Δ = 7.6%; *CXCL11* Δ = 1.7%). Similarly small increases in the MSP-rich fraction were observed after IFNγ rechallenge relative to primary stimulation (*IDO1* Δ = 6.9%; *CXCL11* Δ = 3.8%). These shifts were considerably smaller than the corresponding differences measured by flow cytometry for IDO1 (**Fig. 1B**). Within the resolution of this analysis, these results suggest that accessibility at the promoters of *IDO1* and *CXCL11* largely return to baseline during rest and exhibit no substantial change from the initial stimulation upon rechallenge.

**Fig. 4.**
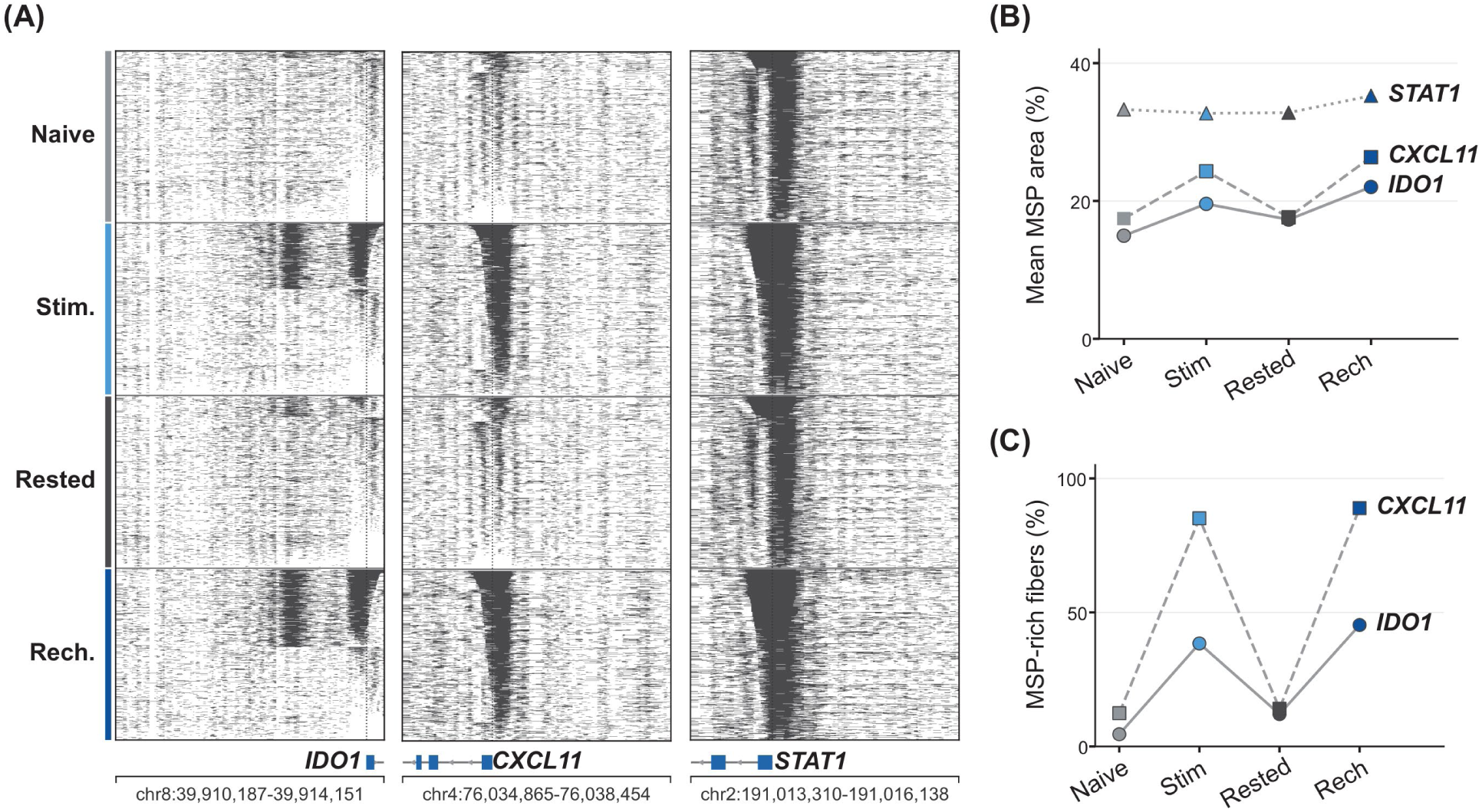
Promoter accessibility tracks stimulation but is not retained between IFNγ exposures: **(A)** Single-fiber DAF-seq modification-sensitive patch (MSP) maps across the IFNγ training regime (Fig. 1A) at the promoter amplicons of the memory genes *IDO1* and *CXCL11* and the non-memory control STAT1; amplicons ∼4.1, 3.7, and 2.9 kb, respectively. Each horizontal line is one DNA fiber; dark shading marks MSP-covered (accessible) chromatin. **(B)** Mean MSP-covered area per fiber across each amplicon versus condition. Series: *STAT1* (triangles), *CXCL11* (squares), IDO1 (circles); marker fill shades by condition. **(C)** Fraction of MSP-rich fibers called by k-means clustering (k = 2) versus condition, for *CXCL11* (squares) and *IDO1* (circles); marker fill shades by condition.

Despite only identifying minor hysteretic changes in accessibility, DAF-seq was able to resolve a memory effect associated with transcription factor binding (**Fig. S9**). Here, the FiberHMM model was used to detect transcription-factor-sized footprints on fibers and then map protein occupancy across individual motifs. With this analysis, TF footprint densities rose in the promoter-proximal window within 150 bp of the *IDO1* TSS and at several promoter-proximal sites in *CXCL11* after stimulation or rechallenge (**Figs. S9A,C**). After either IFNγ exposure, footprint density peaks localized to positions matching two known IFNγ-response motifs, STAT/ISGF3 and IRF1 (**Figs. S9B,D**). Footprint densities in naive and rested cells were nearly indistinguishable, but footprint density was higher in rechallenged than in stimulated cells. Together, these results indicate that stimulus responsive TF occupancy at *IDO1* and *CXCL11* promoters was not maintained through the resting period but was reacquired to a greater extent upon rechallenge. The potentiated transcription of memory genes is therefore likely linked to the elevated TF occupancy at these sites.

### H3K27me3 hysteresis delineates highly-responsive memory genes

To evaluate whether H3K27me3 hysteresis represents a distinct memory signature across the broader set of IFNγ-responsive genes, we profiled H3K27me3 spatial occupancy around the transcriptional start site (TSS) using CUT&RUN sequencing data across all ISGs. We first assessed baseline spatial distributions of H3K27me3 in naive MSCs by comparing mean H3K27me3 signals (Counts Per Million (CPM)) across the three response classes: the 50 memory genes showing potentiated mRNA transcription (log_2_(FC_rechall / stim_)> 0.5), the 30 refractory genes showing the opposite behavior (log2 ratio ≤ −0.5), and the remaining 983 non-memory genes (−0.5 < log2 ratio < 0.5) (**Fig. 5A**). Average traces of H3K27me3 enrichment surrounding the promoters of the 50 memory genes exhibited significantly elevated baseline H3K27me3 levels compared to non-memory ISGs (**Fig. 5A**). Mean CPM values across a ±3 kb window surrounding the TSS were approximately 4.1-fold higher for memory genes than for non-memory ISGs (⟨CPM⟩ = 0.110 and 0.027, respectively; Mann-Whitney P = 7.4 × 10⁻¹¹) (**Fig. S10A**). This difference demonstrates a clear baseline propensity for memory loci to harbor repressive H3K27me3 marks prior to stimulation. Nevertheless, despite this apparent enrichment among highly potentiated memory ISGs, H3K27me3 levels varied appreciably across non-memory ISGs, resulting in significant overlap between the two distributions **(Fig. S10A**). We attribute this overlap to the broad baseline presence of this repressive mark at ISG promoters in naive cells. Thus, while H3K27me3 appears to be selectively enriched in responsive memory ISGs, a significant portion of non-memory ISGs also carry appreciable H3K27me3 marks in their naive state.

**Fig. 5.**
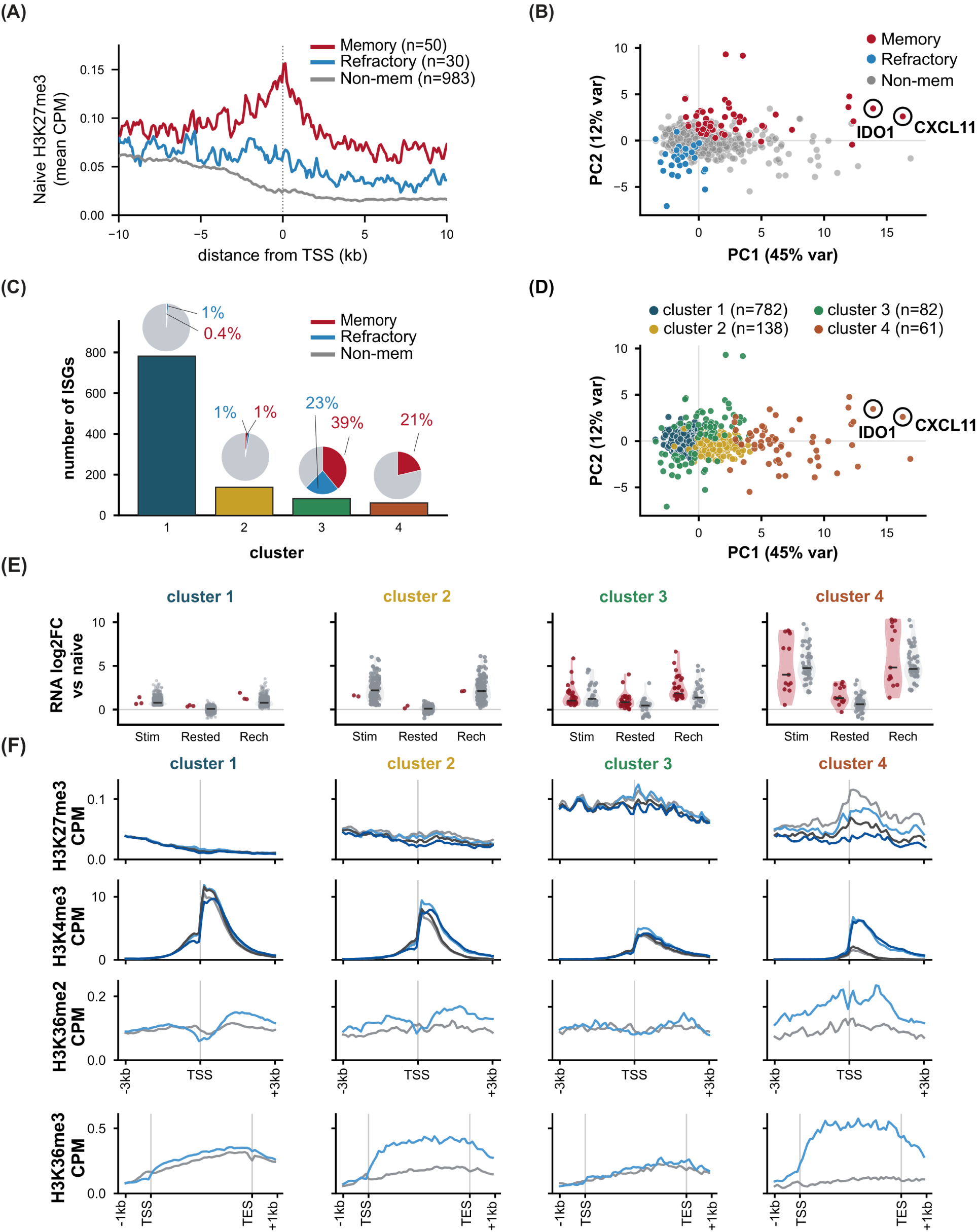
H3K27me3 hysteresis delineates highly-responsive memory genes: **(A)** Aggregate H3K27me3 CUT&RUN signal (mean CPM, Naive) over a ±10 kb window centered on the TSS, averaged separately for memory (n = 50, red), refractory (n = 30, blue), and non-memory (n = 983, grey) **(B)** Multi-omic PCA of IFNγ-responsive genes (PC2 versus PC1), colored by classification: memory (red), refractory (blue), non-memory (grey); *IDO1* and *CXCL11* circled. **(C)** Number of ISGs assigned to each of the four Gaussian-mixture clusters, with pie insets giving the fraction of each cluster classified as memory (red) or refractory (blue). **(D)** The same PCA as in B, colored by GMM cluster assignment (cluster 1, teal; cluster 2, yellow; cluster 3, green; cluster 4, orange); *IDO1* and *CXCL11* circled. **(E)** Per-cluster RNA log2 fold change relative to naive at Stimulated, Rested, and Rechallenged; memory genes in red, non-memory in grey. **(F)** Per-cluster metaplots. Rows, top to bottom: H3K27me3 and H3K4me3 over a TSS-centered ±3 kb window (all four conditions), then promoter H3K36me2 (±3 kb) and gene-body H3K36me3 (scaled TSS→TES with ±1 kb flanks) for Naive and Stim only. Conditions colored per the shared legend (Naive, grey; Stim, light blue; Rested, dark grey; Rechallenged, dark blue).

To more systematically delineate the regulatory signatures distinguishing memory from non-memory ISGs, we conducted an integrated, multi-omic principal component analysis (PCA) (**Fig. 5B, Fig. S10B,C**). This analysis incorporated transcriptomic and epigenomic datasets across all four experimental states: naive, stimulated, rested, and rechallenged. Feature vectors for PCA were built using a standardized dataset comprising mRNA levels together with promoter (TSS ±3 kb) H3K27me3 and H3K4me3 across all four conditions, as well as promoter H3K36me2 and gene-body H3K36me3 measurements from the naive and stimulated conditions. These parameters captured key dynamic transcriptional and epigenetic responses, including primary induction magnitude, persistence through the rest period, and rechallenge potentiation. This integrated framework enabled separation of the core ISG response classes (memory, refractory, and non-memory). Principal Component 1 (PC1) accounted for 45% of the total variance in the data and was weighted heavily by the overall magnitude of transcriptional activation, depletion of H3K27me3 across treatments, potentiation of H3K4me3 upon rechallenge, and accumulation of both H3K36 marks upon stimulation. Memory and refractory loci diverged sharply along PC1, with refractory ISGs enriching toward negative scores and memory ISGs spanning positive values. Highly responsive memory loci (log_2_(FC_rechall / stim_)> 0.5), particularly *IDO1* and *CXCL11*, localized to the highest-magnitude extreme of PC1, which is consistent with their robust IFNγ induction, extensive histone PTM hysteresis, and significant transcriptional potentiation. In contrast, non-memory genes remained tightly clustered near the origin across PC1, PC2, and PC3 despite subtle dispersion.

Using a Gaussian mixture model (GMM) to cluster ISGs based on their multi-omic feature vectors, we partitioned ISGs into four distinct clusters exhibiting different combinations of dynamic transcriptional and chromatin behaviors (**Fig. 5C and 5D, Fig. S11A,B**). Clusters 1 and 2 were primarily composed of non-memory ISGs that exhibited little transcriptional hysteresis (**Fig. 5C**). Cluster 1 contained the vast majority of non-memory ISGs (782 of 1,063 analyzed genes; 73.6% of all ISGs). These loci exhibited modest primary induction and minimal baseline or stimulus-induced changes in chromatin marks. In contrast, cluster 3 contained nearly all of the refractory and a subset of memory-classified genes. The most highly-responsive ISGs, including *IDO1* and *CXCL11*, were grouped into cluster 4, which displayed all of the hallmark memory responses described above including: strong primary transcriptional activation, marked potentiated expression upon rechallenge, and robust induction of activating histone marks (H3K4me3, promoter H3K36me2, and gene-body H3K36me3) (**Fig. 5E and 5F, right column**). Furthermore, in contrast to clusters 1-3, cluster 4 exhibited the most pronounced hysteresis of H3K27me3, characterized by a progressive depletion of this repressive mark upon activation, and through both the rest and rechallenge (**Fig. 5F and Fig. S11C**). Consistent with the above analyses focused on *IDO1* and *CXCL11*, this result further suggests that the persistent loss of H3K27me3 may constitute a core signature of memory gene subsets that exhibit high overall activation and potentiation upon challenge. Nevertheless, cluster 4 still also harbored a significant proportion of non-memory ISGs (13 memory ISGs vs. 48 non-memory ISGs). Focused analysis of this subset shows that these non-memory ISGs also share core signatures of hysteretic H3K27me3 depletion and stimulus-responsive gains in H3K4me3, H3K36me2, and H3K36me3, without displaying transcriptional memory at the mRNA level (**Fig. S11D**). Consequently, while H3K27me3 hysteresis appears central to the establishment of memory at highly stimulus-responsive loci such as *IDO1* and *CXCL11*, additional cis-regulatory signatures and epigenetic features are also likely to have roles in determining whether persistent H3K27me3 depletion manifests in hysteresis at the mRNA transcriptional level.

## DISCUSSION

Following exposure to inflammatory signals or pathogens, cells frequently develop memory towards these stimuli that dictates their responsiveness to future environmental cues. This hysteretic behavior relies on heritable epigenetic modifications that influence secondary transcriptional activation dynamics. As demonstrated here and by others (*6–8*, *12*, *18*, *39*, *40*), memory behaviors are highly locus-specific and variable. Some stimulus-responsive loci return fully to a baseline transcriptional state and then show potentiated secondary expression kinetics, a behavior we refer to as ‘classical memory’. Conversely, other pathway genes become tolerized and display attenuated or refractory behaviors, while still others exhibit complex, multi-phasic responses where elevated transcriptional states persist even during resting periods after the signal is removed. Understanding the regulatory logic that links chromatin remodeling to these diverse outcomes remains challenging. It is still unclear what locus-specific features govern susceptibility to chromatin-based memory, and what ultimately dictates the direction and magnitude of a gene’s hysteretic response. It is also unclear whether the transcriptional machinery itself, such as the physical act of transcription or the localized accumulation of RNA Polymerase II (RNAPII), actively installs these epigenetic bookmarks, or if changes driven by composite activation pathways ultimately dominate the chromatin remodeling process. In this study, we show that primary mesenchymal stromal cells (MSCs) exhibit robust, long-term transcriptional memory following exposure to the pro-inflammatory cytokines IFNγ and TNFα. Although both cytokines induce extensive expression changes and highly diverse hysteretic responses across the MSC genome, the use of selectively targeted CRISPR activation (CRISPRa) as a transient primary activating signal demonstrates that locus-specific initiation of transcription are sufficient to install durable memory entirely independently of these complex signaling cascades.

While locus-specific perturbations via CRISPR inhibition (CRISPRi) and other synthetic tools have successfully characterized how repressive chromatin modifications mediate heritable silencing (*41–43*), efforts to investigate mechanisms of memory using CRISPR activation (CRISPRa) have yielded a range of results. For example, transient transfection and washout of DNA plasmid constructs (dCas9-SAM) was not found to establish memory at clusters of guanylate-binding protein (*GBP*) genes in HeLa cells, even though co-treatment with CRISPRa and IFNγ synergistically elevated ISG expression (*24*). This suggested that locus-specific transcriptional initiation alone is insufficient to induce memory. Conversely, transient activation of Polycomb-repressed genes in immortalized mouse embryonic fibroblasts by dCas9-VPR established a toggle-switch-like memory, where target genes remained active even after the dCas9 complex was diluted (*25*). This toggle switch behavior has been attributed to a positive feedback loop where active transcription sustains continuous expression. Although the underlying mechanisms of memory responses may naturally vary depending on the target gene, CRISPR tool, and cell line, the ability to generate memory at ISGs that return to baseline in rested cells through transient mRNA delivery of VP64-dCas-VP64 demonstrates that memory can be established via local transcriptional activation and maintained without requiring continuous feedback from the transcriptional machinery. Interestingly, this memory persists even though CRISPR-mediated expression levels were lower than those induced by IFNγ, indicating that the initial magnitude of expression is not the primary driver of memory formation. Furthermore, *CXCL11* resides within a cluster of multiple memory genes that typically exhibit a classical transcriptional memory response. The ability to selectively activate *CXCL11* without affecting the neighboring *CXCL9* and *CXCL10* genes suggests that localized memory formation can occur without the coordinated activation of the surrounding cluster.

Local activation via VP64-dCas-VP64 also induced persistent chromatin changes, notably a depletion of H3K27me3 that was maintained in rested cells and progressed further upon restimulation. Given that pharmacological inhibition of H3K27me3 was also found to install memory-like behavior by sensitizing *IDO1* to subsequent IFNγ exposure, this resting-state signal points to H3K27me3 depletion and Polycomb Repressive Complex 2 (PRC2) inhibition as mechanistic drivers of the long-term memory responses at highly transcribed ISGs like *IDO1* and *CXCL11*. While suggesting memory does not stem from increase promoter accessibility, single-molecule analyses via DAF-seq suggests the potentiated transcription of these genes is linked to enhanced TF binding in rechallenge, as seen in prior CUT&RUN-seq experiments (*24*). This indicates the H3K27me3 depletion yields a more permissive chromatin environment for TF engagement. It is also possible that the high induction of H3K36me2/3 reinforces the depletion of H3K27me3 through a positive feedback loop. H3K27me3 is a well-established barrier to transcriptional activation at developmentally regulated genes (*44*, *45*). The transcription-associated post-translational modifications (PTMs) H3K4me3 and H3K36me2/3 can biochemically antagonize H3K27me3 deposition (*46*, *47*), and the accumulation of H3K36me2/3 is linked to H3K27me3 loss in cancer (*48*, *49*) and inflammation (*50*). As RNA Polymerase II (RNAPII) elongates during active transcription, its hyperphosphorylated C-terminal domain (CTD) directly recruits histone methyltransferases to actively deposit H3K36me3 across the transcribed gene body (*22*). Crosstalk between these marks could therefore create a feedback loop where the initial act of transcription itself reinforces the maintenance of a transcriptionally quiescent, but primed state in rested cells without requiring continuous RNAPII activity to maintain memory.

Finally, our multi-omics data reveal that while progressive H3K27me3 depletion is a key feature of transcriptional memory, this chromatin trajectory is not confined exclusively to memory genes. A substantial subset of non-memory ISGs follows a similar trajectory without exhibiting enhanced expression upon rechallenge. This decoupling shows that H3K27me3 hysteresis does not dictate transcriptional outputs independently. Rather, local *cis*-regulatory features likely determine whether this primed epigenetic state is realized as transcriptional memory. Fully interpreting these dynamics is challenging and limited by the significant complexity of histone modification patterns, underscoring the need to quantify the temporal dynamics of the broader spectrum of histone marks. While progressive H3K27me3 depletion has been similarly identified as an epigenetic signature of IFNγ-induced memory in HeLa cells (*18*), this effect was accompanied by additional complex chromatin landscape adaptations marked by the elevation of permissive marks, such as H3K4me1, H3K14ac, and H4K16ac. Elevated H3K27me3 was also observed after initial IFNγ stimulation at GBP cluster genes, a dynamic not seen in our study. Furthermore, H3K36me3 has been observed at memory genes following short-term (48-hour) IFN-β stimulation (*7*). Considering it is highly likely that biochemical antagonisms, kinetic delays in histone turnover, and the recruitment of chromatin remodeling complexes all play coordinated roles, determining the mechanisms that suppress the long-term local reacquisition of H3K27me3 and dictate secondary enhanced transcriptional outputs will require more comprehensive, time-resolved proteomic analysis of chromatin organization. Nevertheless, given the complexity of these stimulatory pathways, the multifaceted hierarchy of epigenetic changes, and dependencies on local chromatin landscapes, locus-specific perturbations via CRISPRa or similar synthetic tools will be critical for resolving these distinctions.

## MATERIALS AND METHODS

### Cell culture and MSC banking

Human adipose-derived mesenchymal stromal cells (MSCs) were obtained from ATCC (Adipose-Derived Mesenchymal Stem Cells, Normal, Human; ATCC PCS-500-011). Experiments used banked working stocks from three donors (lots 80921243, 80705221, and 70040251). Cells were received at passage 2, expanded one passage, and cryopreserved in multiple vials at 1 × 10⁶ cells/ml in 10% DMSO in complete culture medium. Cells were thawed, expanded, and used for experiments at passage 4 to 8; higher passages were avoided. All cultures were maintained at 37 °C and 5% CO₂, seeded at 5,000 cells/cm² for routine expansion, and fed every 2 to 3 days. Cells were dissociated with 0.05% trypsin-(Gibco cat. 25-200-056) EDTA.

MSCs were cultured in low-glucose DMEM (1 g/l glucose with L-glutamine and sodium pyruvate; Corning, cat. MT10014CV) supplemented with 5% fetal bovine serum (FBS; MilliporeSigma, cat. F2442), 5 ng/ml recombinant human FGF2 (R&D Systems, cat. CFBTFGFB020), 5 ng/ml recombinant human EGF (R&D Systems, cat. 236-EG-200), and 1% penicillin/streptomycin (Gibco, cat. 15-140-163).

### IFNγ and TNFα training paradigm

Transcriptional memory was assayed with a four-condition training paradigm. MSCs were thawed, expanded once, and split into trained and untrained lineages. The trained lineage received an initial (priming) stimulation with 50 ng/ml recombinant human IFNγ (STEMCELL Technologies, cat. 50-197-6285) added to the culture medium; 24 h later the cells were washed twice with PBS, re-fed, and subcultured for 10 days to permit mitotic division before rechallenge. At the end of the rest interval, the trained lineage and a matched untrained lineage were restimulated with 50 ng/ml IFNγ for 10 h and then harvested. This design defined four conditions used throughout: Naive (never exposed to IFNγ), Stimulated (untrained cells given a single 10 h IFNγ stimulus before harvest), Rested (trained, rested 10 days, harvested without restimulation), and Rechallenged (trained, rested 10 days, then restimulated 10 h before harvest).

For memory-persistence experiments, the rest interval before rechallenge was varied across 4, 10, and 21 days. For the TNFα variant, the identical paradigm was run with 10 ng/ml recombinant human TNFα (HEK293-expressed, carrier-free; R&D Systems, cat. 10291-TA) for the 24 h priming and 10 h rechallenge, with trained cells restimulated with TNFα before harvest. Cells for molecular readouts (CUT&RUN, DAF-seq, RNA-seq) were stimulated in 15 cm dishes at approximately 1 × 10⁶ cells per dish.

### EZH2 inhibition

To deplete H3K27me3, MSCs were treated with the EZH2 inhibitor tazemetostat (EPZ-6438; APExBIO, cat. A8221) dissolved in DMSO at 1 µM final concentration, with a matched 0.1% DMSO vehicle control. The working concentration was selected from an Alamar Blue viability titration across 0.1 to 10 µM. Inhibitor and medium were refreshed every 2 to 3 days over a 6-day treatment window, after which cells were stimulated with IFNγ for 10 h and harvested for RT-qPCR. Global H3K27me3 depletion was confirmed by western blot against total histone H3 (see Western blotting).

### Cell proliferation and division tracking

Proliferation was quantified by serial cell counting. Cells were stimulated with IFNγ as above, replated at 50,000 cells/well in 6-well plates, and counted by trypan blue exclusion every 72 h across three successive passages, replating 50,000 cells at each interval. Population doubling time and cumulative population doublings were calculated from the counts. Growth rate was computed as µ = (ln N_t − ln N_0)/t; doubling time as ln(2)/µ; and the number of divisions per interval as the per-day growth rate multiplied by the interval length, summed across passages.

### mRNA synthesis for CRISPR activation

CRISPRa effector mRNAs VP64-dCas9-VP64, were produced by in vitro transcription (IVT). Transcription templates were T7-promoter plasmids built with the Takara Cloning Kit for mRNA Template (cat. 6143) the coding sequence for VP64-dCas9-VP64 was amplified from addgene plasmid #113341 (*51*) and cloned by Gibson assembly into the provided Takara T7 backbone, then linearized with HindIII-HF (New England Biolabs, cat. R3104S) before use. IVT was performed with the TriLink CleanCap AG (3’ OMe) CleanScript IVT Kit (cat. K-7413) according to the manufacturer’s specifications, typically in 20 µl reactions, which yielded sufficient mRNA for experiments. mRNA was purified by lithium chloride precipitation, washed in 70% ethanol, resuspended in nuclease-free water, and stored at −80 °C. Product integrity (approximately 4,500 nt for the CRISPRa effectors) was verified on a 1% agarose gel with NEB RNA Loading Dye (2X, cat. B0363S) and an ssRNA ladder (New England Biolabs, cat. N0362S).

### sgRNA design and selection

Single-guide RNAs were chemically synthesized and chemically modified (Synthego, CRISPRevolution sgRNA EZ Kit, 1.5 nmol; 2’-O-methyl analogs at the first three and last bases, with 3’ phosphorothioate bonds between the first three and last two bases). The *IDO1* guide (protospacer 5’-GTGACCACAGTTTATCACAG-3’, “sg5”) and the *CXCL11* guide (protospacer 5’-AAGCACTGTGAAAACCCAGA-3’) were each selected as the best-performing guide for their locus from the genome-wide CRISPRa library of Sanson et al. (Calabrese set A). (*52*) The *CD34* benchmark guide (protospacer 5’-GAAAGCTGAACGAGGCATC-3’; chr1:207,911,360-207,911,378) was taken from Johnston et al. (*53*) A Synthego Negative Control (scramble) sgRNA served as the non-targeting control (protospacer 5’-GCACTACCAGAGCTAACTCA-3’).

### CRISPRa mRNA delivery and priming

CRISPRa components were delivered by reverse transfection with Lipofectamine MessengerMAX (Invitrogen, cat. LMRNA015) at 1.5 µl per ml of transfection medium. Effector mRNA and sgRNA were combined at a 4:1 mass ratio (1 µg VP64-dCas9-VP64 mRNA and 0.25 µg sgRNA per ml of transfection medium) in Opti-MEM (Gibco, cat. 31985070); the mRNA/sgRNA mix and the diluted MessengerMAX were each prepared in Opti-MEM, combined, incubated 10 min and then 5 min at room temperature, and dispensed into the vessel before MSCs were added on top. Medium was fully changed 16 to 24 h after transfection. Transfections were scaled by culture volume, up to 15 cm dishes for genomics-scale experiments. For synthetic-memory (priming) experiments, cells were transfected on day 0 with an *IDO1*-targeting, *CXCL11*-targeting, or non-targeting sgRNA, rested in culture for 10 days, and then given their first 50 ng/ml IFNγ stimulation for 10 h before analysis by flow cytometry, western blot, RT-qPCR, CUT&RUN-qPCR, or RNA sequencing.

### Flow cytometry

Intracellular IDO1 was quantified by flow cytometry. Cells were lifted with 0.05% trypsin-EDTA, counted, washed in staining buffer (1× PBS, 1% FBS, 0.09% sodium azide), fixed and permeabilized in BD Cytofix/Cytoperm (BD Biosciences, cat. 554714) for 20 min at 4 °C, and washed in 1× BD Perm/Wash. Cells were stained (approximately 1:10 in Perm/Wash, 30 min at 4 °C) with an anti-IDO1 antibody, either the FITC conjugate (clone eyedio; eBioscience, cat. 11-9477-42) or the PE conjugate (cat. 12-9477-42), then washed and resuspended in staining buffer. Data were acquired on a Sony SA3800 spectral analyzer or a Sony MA900 and analyzed in FlowJo (V10.10.2). Surface CD34 was stained without permeabilization using Alexa Fluor 647 anti-human CD34 (BioLegend, cat. 50-207-6593, via Fisher) for 30 min at room temperature.

### Western blotting

Whole-cell lysates were prepared in RIPA buffer (Thermo, cat. 89900) with Halt Protease Inhibitor Cocktail (100×; Thermo, cat. 78430), clarified at 14,000 rpm for 15 min at 4 °C, and quantified by BCA assay (Thermo, cat. 22327; read on a BMG CLARIOstar plate reader). For histone-modification blots (H3K27me3 and total H3), histones were acid-extracted: nuclei were isolated in Triton extraction buffer (PBS with 0.5% Triton X-100, 2% protease inhibitor, 0.02% sodium azide, and 5 mM sodium butyrate) and pelleted at 650 × g for 10 min at 4 °C, histones were extracted in 0.2 N HCl overnight at 4 °C, and the extract was neutralized with 2 M NaOH (one-tenth volume) following a simplified acid-extraction protocol (Abcam). Equal protein (5 to 25 µg; 1 µg for histones) was denatured in Laemmli buffer with β-mercaptoethanol at 95 °C for 10 min (65 °C for histone samples), resolved on 7.5% Mini-PROTEAN TGX precast gels (Bio-Rad, cat. 4561023) or 4 to 20% gradient gels for small proteins, and transferred to PVDF membrane (Bio-Rad Trans-Blot Turbo RTA Mini LF PVDF kit, cat. 1704274) with a Bio-Rad Trans-Blot Turbo system. Membranes were blocked in EveryBlot Blocking Buffer (Bio-Rad, cat. 12010020), probed overnight at 4 °C with primary antibodies (typically 1:1,000), incubated with HRP-conjugated secondary antibodies (1:2,000; anti-rabbit IgG, Cell Signaling Technology cat. 7074; anti-mouse IgG, Cell Signaling Technology cat. 7076) for 1 h at room temperature, and developed with ECL substrate (Bio-Rad Clarity, cat. 1705061, or Thermo SuperSignal West Femto, cat. 34094) on a Bio-Rad ChemiDoc. β-Tubulin was imaged directly as a loading control using hFAB Rhodamine Anti-Tubulin (Bio-Rad, cat. 12004165).

Primary antibodies were: IDO1 (clone V1NC3IDO; eBioscience/Invitrogen, cat. 14-9750-82), phospho-STAT1 (Y701) (Invitrogen, cat. MA5-15071), total STAT1 (Proteintech, cat. 66545-1-Ig), IRF1 (Abcam, cat. ab243895, clone BLR039F), Cas9/dCas9 (Diagenode, cat. NC1554064), CXCL11 (R&D Systems, cat. MAB672SP; 1:1,000), H3K27me3 (Millipore, cat. ABE44), and total histone H3 (Abcam, cat. ab1791). For CXCL11, cells were stimulated with IFNγ for 24 h with BD GolgiPlug (brefeldin A; BD, cat. 555029) added for the final 6 h to allow intracellular accumulation. Band intensities were quantified in ImageJ.

### RNA extraction and RT-qPCR

Total RNA was extracted with the RNeasy Plus Mini Kit (Qiagen, cat. 74136), with genomic DNA removed on the kit’s gDNA-eliminator column, and quantified on a NanoDrop OneC. cDNA was synthesized from 250 ng RNA with the iScript Advanced cDNA Synthesis Kit (Bio-Rad, cat. 1725038BUN) in a 10 µl reaction and diluted 1:10. Quantitative PCR was performed in 10 µl reactions containing 5 µl Luna Universal qPCR Master Mix (New England Biolabs, cat. M3003E), 0.5 µl primer mix, and 4.5 µl diluted cDNA on a Bio-Rad CFX96 Touch Deep Well real-time system (cat. 3600037; CFX Maestro v5.2.008.0222), with cycling of 95 °C for 60 s, then 40 cycles of 95 °C for 15 s and 60 °C for 30 s, followed by a melt curve. Expression was normalized to *ACTB* and quantified by the 2^(−ΔΔCt) method; primer sequences are listed in table S1.

### RNA sequencing

Three bulk RNA-seq datasets were generated: an IFNγ training set (four conditions, n = 3), a TNFα set (n = 2), and a CRISPRa *CXCL11* synthetic-memory set (*CXCL11*-targeting versus non-targeting cells, rested or restimulated with IFNγ at 10 and 21 days of rest, n = 3). Total RNA was extracted with the RNeasy Plus Mini Kit (Qiagen), quantified on a Qubit 4.0 fluorometer, and assessed for integrity on an Agilent TapeStation 4200 (RIN 9.6 to 10.0 for the IFNγ set). For the IFNγ and TNFα sets, libraries were prepared by Azenta/GENEWIZ with the NEBNext Ultra II RNA Library Prep Kit for Illumina using Oligo-d(T) poly-A enrichment, and sequenced on an Illumina NovaSeq in a 2 × 150 bp paired-end configuration targeting approximately 20 million reads per sample; base calls were demultiplexed with bcl2fastq 2.20. The CRISPRa CXCL11 set was library-prepped and sequenced by Plasmidsaurus, which also performed primary read processing and gene-level quantification (see RNA-seq data analysis). Libraries carried unique molecular identifiers and yielded 15.9 to 20.6 million reads per sample.

### RNA-seq data analysis

The IFNγ and TNFα training sets were processed with a single pipeline. Reads were aligned and quantified with RSEM (v1.3.3) against a STAR (v2.7.11b) index of the human genome (hg38, GENCODE v49 annotation), and per-gene expected counts were imported into R with tximport (v1.36.0; type = “rsem”). Differential expression was computed with DESeq2 (v1.50.2; R v4.5.1) using design ∼ condition, after removing genes with fewer than 10 summed counts across samples. Variance-stabilized counts were used for principal component analysis (DESeq2 plotPCA on the top 500 most variable genes), and volcano and MA plots applied lfcShrink with thresholds of adjusted P < 0.05 and |log2FC| > 0.5.

The CRISPRa CXCL11 synthetic-memory set was pre-processed by Plasmidsaurus on a different pipeline and delivered as a gene-level count matrix. Reads were trimmed with fastp (v0.24.0; poly-X and 3’ quality trimming, minimum Phred 15, minimum length 50 bp), aligned with STAR (v2.7.11) with non-canonical splice junctions removed, coordinate-sorted with samtools (v1.22.1), and deduplicated on unique molecular identifiers with UMIcollapse (v1.1.0); across the 24 libraries, 15.9 to 20.6 million input reads per sample gave 49 to 59% retention after deduplication. Gene-level counts were generated with featureCounts (subread v2.1.1) using strand-specific counting and fractional assignment of multi-mapping reads, with exons and 3’ UTRs as feature identifiers grouped by gene_id. These counts were rounded to integers and analyzed with DESeq2 (pydeseq2) independently for each timepoint, using a single four-level condition factor (non-targeting or CXCL11-targeting sgRNA, rested or IFNγ-restimulated) and the same count filter and significance thresholds as above.

### Gene set enrichment analysis

Gene set enrichment analysis was performed with clusterProfiler (v4.12.6) gseGO under R 4.4.1 and Bioconductor 3.19. For each contrast against naive cells, genes were ranked in decreasing order by the DESeq2 Wald statistic after removing genes with an undefined statistic, so that a positive normalized enrichment score (NES) indicates enrichment in the stimulated or rested condition. Gene Ontology Biological Process terms were taken from org.Hs.eg.db (v3.19.1) and GO.db (v3.19.1; GO release 2024-01-17) matched on gene symbols, and restricted to sets of 10 to 500 genes. Significance was estimated with the adaptive multilevel permutation procedure of fgsea (v1.30.0), which clusterProfiler invokes by default and which floors reported p values at 1 × 10⁻¹⁰, and p values were adjusted across all tested gene sets by the Benjamini-Hochberg method. Terms with an adjusted p value < 0.05 were considered enriched.

### ChIP-qPCR for dCas9 occupancy

Occupancy of the CRISPRa effector at the *IDO1* promoter was measured by ChIP-qPCR at 1 and 10 days after transfection. Cells were crosslinked in 1% formaldehyde for 5 min at room temperature and quenched in 0.125 M glycine. Nuclei were isolated in Farnham lysis buffer (5 mM PIPES pH 8.0, 85 mM KCl, 0.5% NP-40, protease inhibitor), and chromatin was sonicated in RIPA sonication buffer (1× PBS, 1% NP-40, 0.5% sodium deoxycholate, 0.1% SDS) on a Bioruptor Pico (Diagenode) for 15 cycles of 30 s on/30 s off to a target size of 100 to 500 bp. Pooled chromatin was cleared, 1% was retained as input, and the remainder was immunoprecipitated overnight at 4 °C with 5 µg anti-Cas9 antibody (Diagenode, cat. C15200216-100, clone 4G10; Fisher NC1554064) or 5 µg matched IgG control (Invitrogen, cat. A10630), captured on Protein G Dynabeads (Thermo Fisher, cat. 10003D), washed through low-salt, high-salt, LiCl, and TE buffers, eluted, reverse-crosslinked, and purified on a Qiagen MinElute column (cat. 28004). qPCR was performed in technical triplicate with Luna Universal qPCR Master Mix on a Bio-Rad CFX real-time system at the sgRNA target site and a distal negative-control locus, and occupancy was expressed as percent input. Three biological replicates were analyzed per condition. Primer sequences are listed in table S1.

### CUT&RUN

CUT&RUN was performed with the CUTANA ChIC/CUT&RUN Kit (EpiCypher, cat. 14-1048). Nuclei were extracted before the assay to increase DNA yield: cells were dissociated, pelleted, resuspended in EpiCypher Nuclei Extraction Buffer (cat. 21-1026), incubated on ice, and pelleted, and 500,000 nuclei were bound to activated Concanavalin A beads (11 µl per reaction). A SNAP-CUTANA K-MetStat spike-in panel (EpiCypher, 1 µl per reaction) was included for optional normalization and antibody-target validation. Antibody was bound, pAG-MNase (supplied with the kit) was activated with calcium, and released chromatin was purified and prepared into libraries with the CUTANA CUT&RUN Library Prep Kit (EpiCypher, cat. 14-1002). Libraries were quality-checked on a TapeStation, pooled equimolar, and sequenced 2 × 150 bp on an Illumina NovaSeq X Plus (Admera Health). H3K4me3 and H3K27me3 were profiled across all four training conditions, and H3K36me2 and H3K36me3 in the Naive and Stimulated conditions. Because focal changes in the abundant H3K27me3 mark are read-sampling-limited at standard depth, the eight H3K27me3 libraries were additionally sequenced to high depth (see below). SNAP-certified CUT&RUN antibodies (EpiCypher) were used throughout: H3K4me3 (cat. 13-0060), H3K27me3 (cat. 13-0055), H3K36me2 (cat. RD130056), and H3K36me3 (cat. 13-0058). Antibody binding, K-MetStat spike-in, pAG-MNase targeted digestion, and library amplification followed the manufacturer’s CUTANA kit protocol.

### CUT&RUN data analysis

Reads were trimmed with fastp (v0.24.0; --detect_adapter_for_pe -q 20 -l 25 --cut_right -- cut_mean_quality 15), cropped to 50 bp, and aligned to hg38 with Bowtie2 (v2.5.4; --local --very-sensitive-local --no-unal --no-mixed --no-discordant -I 10 -X 700). ENCODE hg38 blacklist v2 regions were removed with bedtools (v2.31.1). CPM-normalized coverage tracks were generated with deeptools (v3.5.5) bamCoverage (binSize 10, smoothLength 50, extendReads, --ignoreDuplicates), and replicate BAMs were merged for display. Promoter signal was quantified as mean CPM over a TSS ± 3 kb window per gene, and TSS metaplots were computed with deeptools computeMatrix/plotProfile (reference-point TSS, ± 3 kb or ± 10 kb, 100 bp bins) over GENCODE v49 gene models.

To resolve focal H3K27me3, the eight H3K27me3 libraries were concatenated with the original reads and re-sequenced to approximately 118 to 128 million merged paired-end reads per condition (about 10-fold the initial depth) and reprocessed through the same pipeline. A depth-titration analysis subsampled the deep Naive data across a ladder of 10, 20, 40, 60, 80, and 120 million read pairs (five random draws each) and quantified reads in each gene’s TSS ± 3 kb window, with detectability expressed as a Poisson metric (Δreads/√Σreads).

Genome-browser tracks were rendered from replicate-averaged CPM bigWigs with a custom Python tool (pyBigWig/pybigtools), with gene models retrieved from the UCSC API.

### CUT&RUN-qPCR

H3K27me3 enrichment at memory-gene and non-memory-gene promoters was measured by qPCR on CUT&RUN DNA, both across the IFNγ training conditions and after CRISPRa targeting of *IDO1*. CUT&RUN was performed as above with the EpiCypher kit and an H3K27me3 antibody (EpiCypher, cat. 13-0055) in biological triplicate, and purified DNA was quantified by Qubit and diluted to load approximately 0.5 ng per reaction. qPCR was performed in 10 µl reactions with Luna Universal qPCR Master Mix (New England Biolabs, cat. M3003) on a Bio-Rad CFX system (95 °C for 60 s; 40 cycles of 95 °C for 15 s and 60 °C for 30 s; melt curve). Enrichment was expressed as fold change (2^(−ΔΔCt)) relative to a *HOX* control locus and normalized to the Naive (or non-targeting) condition. Primers targeted the *IDO1* and *CXCL11* promoters (within 300 bp of the TSS) with a *HOX* control pair; sequences are in table S1.

### DAF-seq

Chromatin accessibility and transcription-factor occupancy at the *IDO1*, *CXCL11*, and *STAT1* promoters were profiled by DAF-seq (deaminase-assisted single-molecule fiber sequencing). Approximately 1 × 10⁶ MSCs per condition were harvested, and nuclei were isolated in EpiCypher nuclei extraction buffer, washed in isotonic wash buffer (20 mM Tris-HCl pH 7.4, 150 mM NaCl, 3 mM MgCl₂, 10 nM ZnCl₂), and resuspended in 50 µl of the same buffer. Accessible cytosines were deaminated by adding 2 µl of 100 µM SsDddA (a single-stranded DNA cytidine deaminase provided by the Stergachis laboratory) and incubating at 25 °C for 10 min; the reaction was quenched to 5% SDS and genomic DNA was extracted with a Qiagen DNeasy Blood & Tissue kit (cat. 69506). One deaminated reaction per condition per locus was profiled, alongside a non-deaminated control. Promoter amplicons were generated with repliQa HiFi ToughMix (Quanta Bio, cat. 95200) from 1 ng/µl input DNA: *IDO1* (4,064 bp, chr8:39,910,187-39,914,251; primers oGB285/oGB309), *CXCL11* (3,706 bp, chr4:76,034,865-76,038,554; primers oDY002/oDY007), and *STAT1* (2,928 bp, chr2:191,013,310-191,016,238; primers oGB294/oGB303). Primers were designed with degenerate G-to-R substitutions to reduce G-runs and improve deamination readout (IDT). Sequences are in table S2. Amplicons were purified (Qiagen column, or gel extraction where needed) and sequenced on Oxford Nanopore, targeting approximately 5,000 reads per sample. One deaminated reaction per condition per locus was profiled, alongside a non-deaminated control.

### DAF-seq data analysis

Amplicon reads were aligned to hg38 with minimap2 (v2.28) and deamination-decorated with fibertools-rs (v0.7.0) in the upstream DAF-seq processing project, producing per-fiber mapped BAMs with strand tags. Footprints were called on the mapped BAMs with FiberHMM (v2.10.5; fiberseq/Stergachis laboratory) in DddA mode (fiberhmm-call -- enzyme ddda with the transcription-factor recall pass), and modification-sensitive patches (MSPs), nucleosomes, and TF-sized footprints were exported as BED12 (fiberhmm-extract). For each locus and condition, MSP intervals were mapped onto 10 bp bins to build a per-fiber binary accessibility matrix, using fibers covering at least 60% of the plotted interval; conditions were subsampled to equal fiber depth per locus. Per-fiber MSP-covered area was the mean of the binary row, and MSP frequency at the TSS was computed over a TSS ± 150 to 250 bp window. Fibers at the *IDO1* and *CXCL11* promoters were classified as MSP-rich or MSP-poor with a single balanced k-means model per locus (scikit-learn v1.6.1; k = 2, n_init = 20, random_state = 42) fit across all four conditions; *STAT1* was not clustered.

Transcription-factor footprints were analyzed from the FiberHMM TF-sized calls. Amplicon-wide TF density was aggregated into 50 bp bins as the fraction of spanning reads carrying a TF-sized call, with a 150 bp guard band trimmed from each amplicon end. Per-base TF-call density over the TSS ± 100 bp window used only fibers fully spanning the window and was smoothed with a Gaussian kernel (sigma = 2 bp). Sequences in the TSS ± 100 bp window were scanned against JASPAR 2024 CORE vertebrate motifs (retained at ≥ 0.85 of the motif score range), and per-motif footprint occupancy was compared between conditions with one-sided Fisher exact tests and Benjamini-Hochberg correction. Reported motifs (STAT/ISGF3, IRF) are curated. Per-fiber MSP-area comparisons between conditions used two-sided Mann-Whitney U tests with Benjamini-Hochberg correction across the pairwise contrasts, with Cliff’s delta as the effect size. Analyses ran under Python 3.13 with matplotlib 3.10.8, numpy v2.2.3, scipy v1.15.2, pysam v0.23.0, and scikit-learn v1.6.1.

### Multi-omic integration

To partition IFNγ-responsive genes by their transcriptional and chromatin behavior, the 1,063-ISG universe was clustered on 11 per-gene features; all 1,063 genes had complete features and were retained. The features were, all as log2 fold change relative to Naive except the priming contrast: RNA induction (Stimulated), RNA persistence (Rested), and RNA priming (Rechallenged versus Stimulated); promoter (TSS ± 3 kb) H3K27me3 and H3K4me3 at Stimulated, Rested, and Rechallenged; promoter H3K36me2 (Stimulated); and gene-body H3K36me3 (Stimulated). Features were z-scored per feature (correlation PCA), reduced by principal component analysis to three components (scikit-learn v1.6.1 PCA, random_state = 42; PC1 45.4%, PC2 12.2%, PC3 11.2% of variance), and clustered with a Gaussian mixture model (scikit-learn GaussianMixture, covariance_type = “full”, n_init = 10, random_state = 42). The cluster number was selected by the Bayesian information criterion over k = 2 to 10, giving k = 4. Per-cluster memory enrichment was tested with the hypergeometric distribution, and per-cluster chromatin metaplots were computed from replicate-averaged CPM bigWigs (read with pybigtools v0.2.4; promoter marks over TSS ± 3 kb in 100 bp bins; H3K36me3 as a scaled gene-body profile). Analyses ran under Python 3.12 with scipy v1.15.2, numpy v2.2.3, and pandas v2.2.3.

### Statistical analysis

Bar-graph statistics were computed in Python (scipy.stats and statsmodels) and differential-expression statistics with DESeq2, from a single shared analysis module so that all panels follow one set of conventions. Unless stated otherwise, n denotes independent biological replicates: n = 3 for flow cytometry, RT-qPCR, ChIP-qPCR, CUT&RUN-qPCR, and RNA-seq; n = 2 for the CUT&RUN CPM quantifications. Bar graphs show the group mean with all individual replicate values overlaid as points, and error bars are the standard error of the mean (SEM). The statistical test was fixed by experimental design and applied identically wherever that design recurs: two groups were compared by Welch’s unpaired (unequal-variance) t test; the four training conditions (Naive, Stimulated, Rested, Rechallenged) by one-way ANOVA with Tukey’s all-versus-all post-test; multiple guides or conditions against one shared control by one-way ANOVA with Dunnett’s post-test; and the priming/memory design (a 2×2 of guide or treatment by IFNγ) by two-way ANOVA, reporting the interaction term as the priming/memory test, with qPCR tests run on ΔCt values. For RNA-seq expression bars, the bar height is transcripts per million (TPM) and significance is the gene-level DESeq2 adjusted P value (Wald test with Benjamini-Hochberg FDR) for the relevant contrast; no separate test was computed on the TPM values. Genomic proportions (the fraction of genes losing a mark) were compared by Fisher’s exact test, and single-molecule and gene-set comparisons by Mann-Whitney U and paired Wilcoxon signed-rank tests, with Benjamini-Hochberg correction across multiple comparisons; metagene profiles show mean ± SEM. Significance is annotated throughout on the four-tier convention (* P < 0.05, ** P < 0.01, *** P < 0.001, **** P < 0.0001; ns, not significant). The exact test, replicate number, and quantity plotted are given in each figure legend.

## Acknowledgments

We thank Keith Maier (EpiCypher) for early access to the CUTANA Cloud analysis platform, which we used for comparative analysis of our CUT&RUN data. We thank Andrew B. Stergachis and Devaraja Mudeppa (University of Washington) for the kind gift of the SsDddA single-stranded DNA cytidine deaminase that made the targeted DAF-seq experiments possible, and for their guidance on amplicon primer design and validation and on processing the resulting DAF-seq data.

## Author contributions

Conceptualization: G.C.B., I.B.H., M.R.D.

Methodology: M.A.B., G.C.B., S.H.

Investigation: G.C.B., S.H., D.L.Y., D.D., H.S.-B.

Visualization: G.C.B., S.H., D.L.Y., D.D., H.S.-B., M.A.B.

Funding acquisition: G.C.B., I.B.H., M.R.D.

Writing, review and editing: G.C.B., A.R.D., I.B.H., M.R.D.

## Data and materials availability

All data needed to evaluate the conclusions in the paper are present in the paper and/or the Supplementary Materials. All high-throughput sequencing data generated in this study have been deposited in the NCBI Gene Expression Omnibus (GEO) under the following accessions: GSE348230 (bulk RNA-seq of MSCs across the IFNγ and TNFα training paradigms: naive, stimulated, rested and rechallenged, n = 3 and n = 2 biological replicates respectively); GSE347785 (bulk RNA-seq of CRISPRa-primed MSCs, CXCL11-targeting versus non-targeting sgRNA, rested or IFNγ-restimulated at 10 and 21 days of rest, n = 3); GSE347816 (CUT&RUN profiling of H3K4me3 and H3K27me3 across all four training conditions and of H3K36me2 and H3K36me3 in the naive and stimulated conditions, with pooled IgG negative controls, n = 2); and GSE347784 (targeted DAF-seq of the IDO1, CXCL11 and STAT1 promoters across all four training conditions).

All code used to process the sequencing data and to generate the figures and analyses in this paper has been archived at Zenodo and is available at https://doi.org/10.5281/zenodo.22904458. That archived version is the one used for the analyses reported here. The same code is additionally available as a GitHub repository at https://github.com/guybedford98/bedford-2026-ifng-memory. Upstream DAF-seq read processing used the DAF-QC-SMK workflow and FiberHMM, which are distributed separately by the Stergachis laboratory (University of Washington).

This study did not generate new cell lines. Human adipose-derived mesenchymal stromal cells were obtained commercially from ATCC (PCS-500-011), and all antibodies, kits and reagents are commercially available and listed with supplier and catalog numbers in table S3. The SsDddA single-stranded DNA cytidine deaminase used for DAF-seq was a gift from A. B. Stergachis and D. Mudeppa (University of Washington) and is not commercially available; it can be obtained from them under a standard material transfer agreement. Plasmid templates for in vitro transcription of the VP64-dCas9-VP64 and dCas9-eGFP mRNAs were constructed in this study and are available from the corresponding authors on request.

