## Supplementary material for "Induction of hysteretic chromatin states by CRISPRa reveals locus-specific encoding of transcriptional memory": supplumental figures

Guy C. Bedford et al.

**This PDF file includes:**

Figs. S1 to S11
Tables S1 to S3

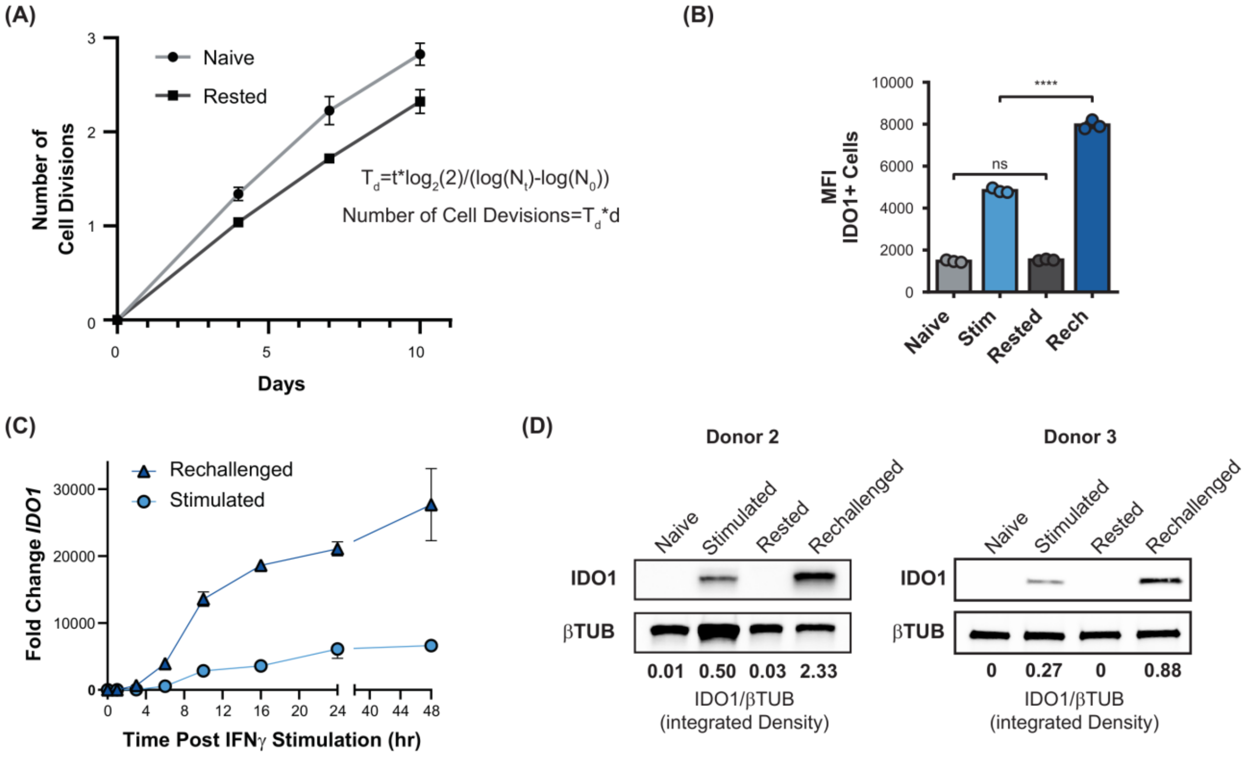


**Fig. S1. *IDO1* inflammatory transcriptional memory is reproducible across donors and shows accelerated, amplified induction on rechallenge.**
**(A)** Cumulative cell divisions of MSCs during the training regime for Naive (grey circles) and Rested (black squares); points, mean; error bars, spread across replicates. Divisions were derived from cell counts using the equations inset on the plot. **(B)** Flow-cytometry MFI within the gated IDO1-positive population across the four training conditions, individual replicates overlaid. **(C)** Time course of *IDO1* transcript induction after IFNγ stimulation for Rechallenged (dark blue triangles) and first-time Stimulated (light blue circles). **(D)** Western blot validation of *IDO1* memory in two additional donor lines; IDO1 and β-tubulin (βTUB) across Naive, Stimulated, Rested, and Rechallenged, with IDO1/βTUB integrated-density ratios printed below each lane. In B, bars are mean ± SEM with individual replicates (n = 3 biological replicates); significance by one-way ANOVA with Tukey's post-test. **** $P < 0.0001$, *** $P < 0.001$, ** $P < 0.01$, * $P < 0.05$; ns, not significant.

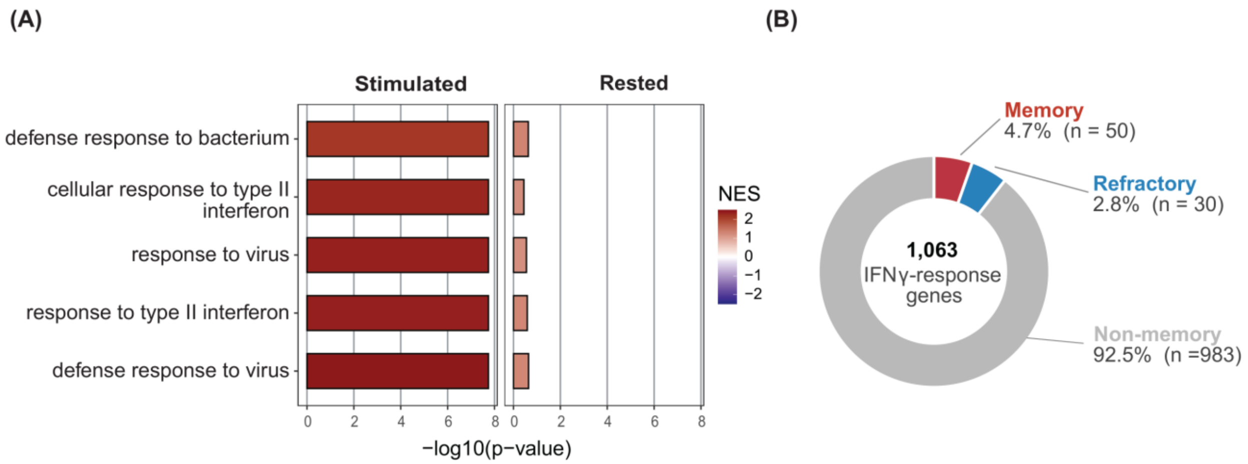


**Fig. S2. IFNγ-response pathway activation resolves after rest.**
**(A)** Gene-set enrichment of the top five Gene Ontology terms, (Stimulated versus Naïve, left; Rested versus Naïve, right). X-axis, −log10(p-value); bar fill, normalized enrichment score (NES). **(B)** Donut chart classifying the 1,063 IFNγ-response genes by rechallenge behavior: Memory (n = 50, red), Refractory (n = 30, blue), Non-memory (n = 983, grey).

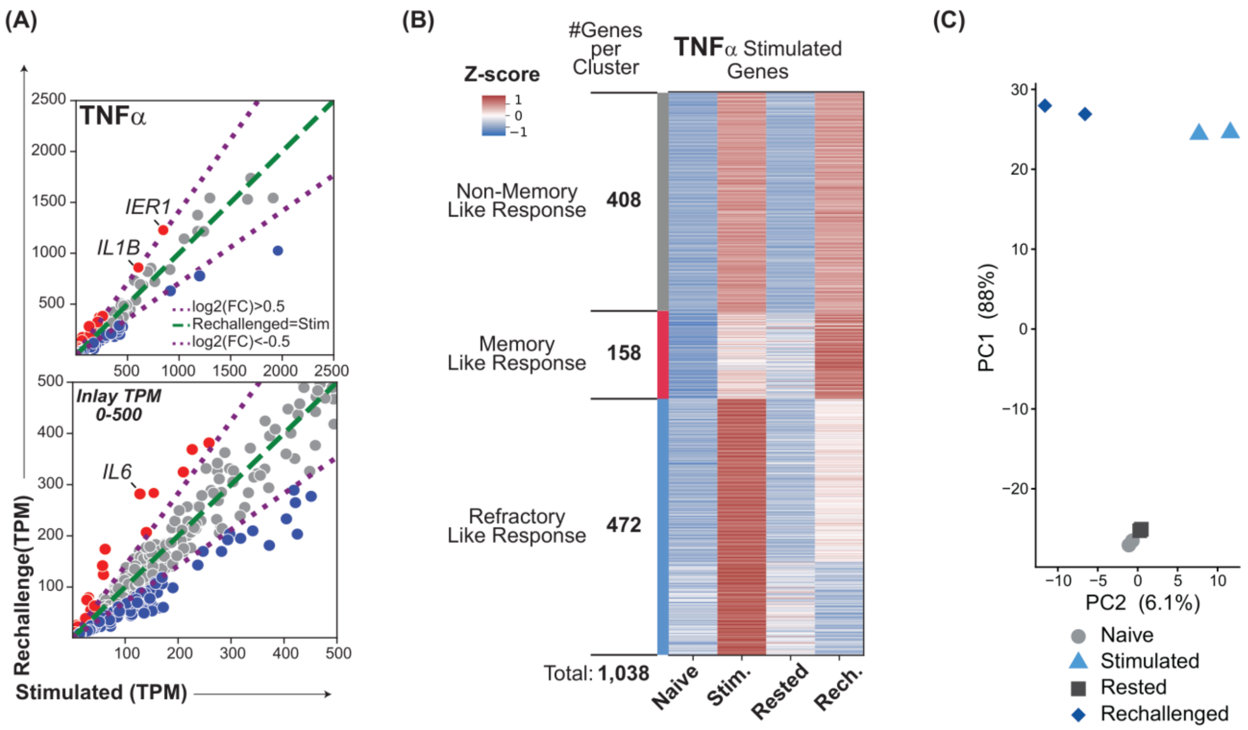


**Fig. S3. TNFα training in MSCs elicits diverse rechallenge behaviors dominated by a refractory response.**

**(A)** Per-gene scatter comparing Rechallenged versus first-time Stimulated expression (TPM) under TNFα training; top, full range; bottom, inset magnifying the 0–500 TPM window. Green dashed line, Rechallenged = Stimulated; purple dotted lines, log2(fold change) = ±0.5. Red points, enhanced on rechallenge; blue, reduced; grey, within threshold. Representative genes labeled. **(B)** K-means heat map (k = 4) of TNFα-stimulated genes (rows) across the four conditions (columns: Naive, Stimulated, Rested, Rechallenged), colored by row-wise z-scored expression, grouped into response classes with per-cluster gene counts at left (total, 1,038 genes). **(C)** Principal-component analysis of the transcriptomes (PC1, 88%; PC2, 6.1%); symbols by condition (Naive, grey circles; Stimulated, light-blue triangles; Rested, dark squares; Rechallenged, blue diamonds), n=2.

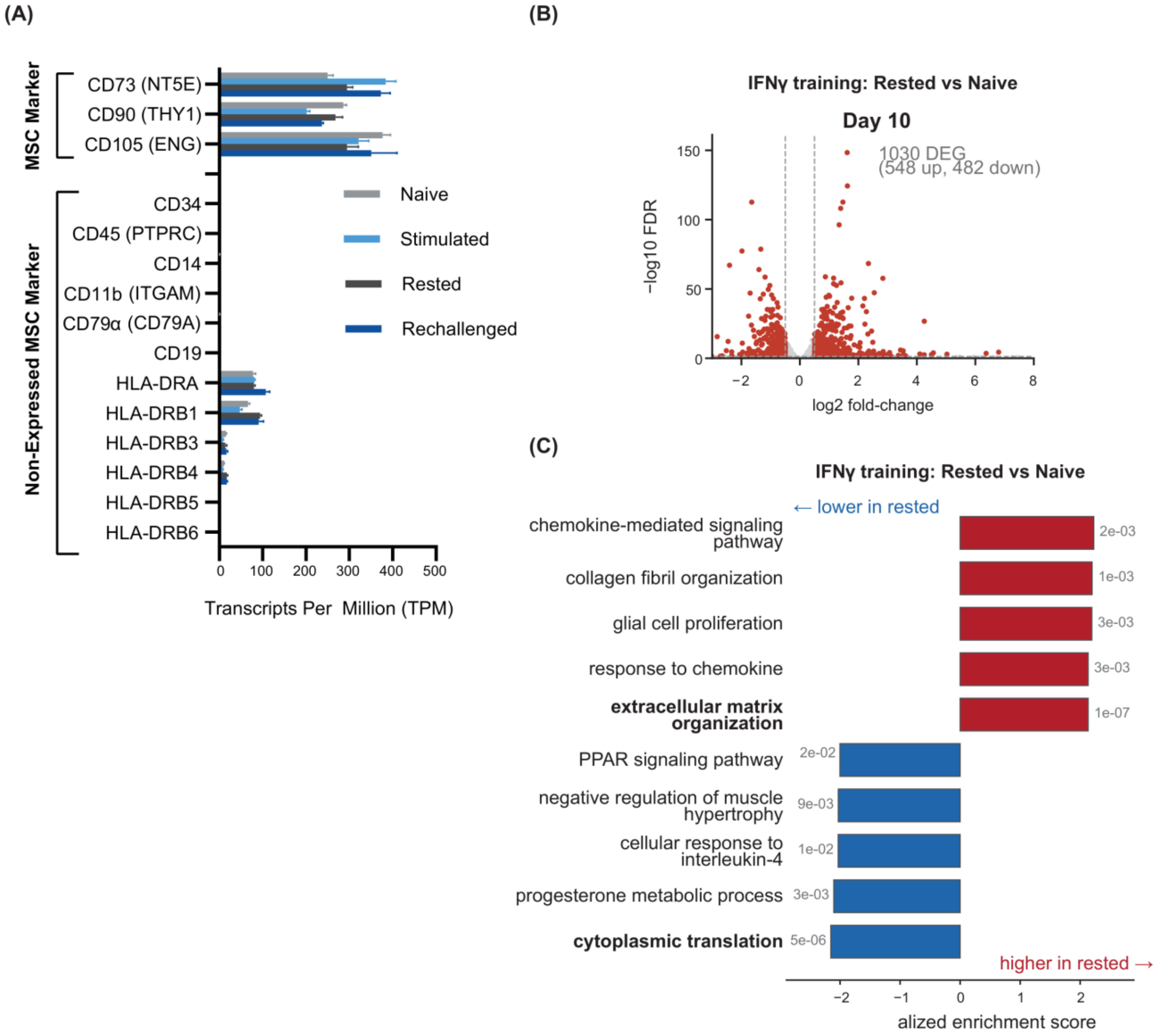


**Fig. S4. Transcriptional drift is apparent 10 days after IFNγ stimulation.**
**(A)** RNA-seq expression (TPM) of International Society for Cellular Therapy MSC identity markers across the naïve stimulated and rechallenged conditions. Top, positive markers (CD73/*NT5E*, CD90/*THY1*, CD105/*ENG*); bottom, markers expected to be absent (*CD34*, CD45/*PTPRC*, *CD14*, CD11b/*ITGAM*, CD79α/*CD79A*, *CD19*, *HLA-DRA/-DRB1/-DRB3/-DRB4/-DRB5/-DRB6*). Bars, mean ± SEM (n = 3 biological replicates). **(B)** Volcano plot of differential expression between the Rested and Naïve conditions at Day 10 of IFNγ training (log2 fold change, Rested/Naïve, versus −log10 FDR); 1,030 DEGs (548 up, 482 down). Significant DEGs in red, non-significant in grey; dashed lines, |log2FC| = 0.5 and FDR < 0.05. **(C)** Gene-set enrichment of the naïve-versus-rested transcriptional drift: normalized enrichment score for the most enriched (red, higher in rested) and most depleted (blue, lower in rested) Gene Ontology biological processes, with the FDR printed beside each bar.

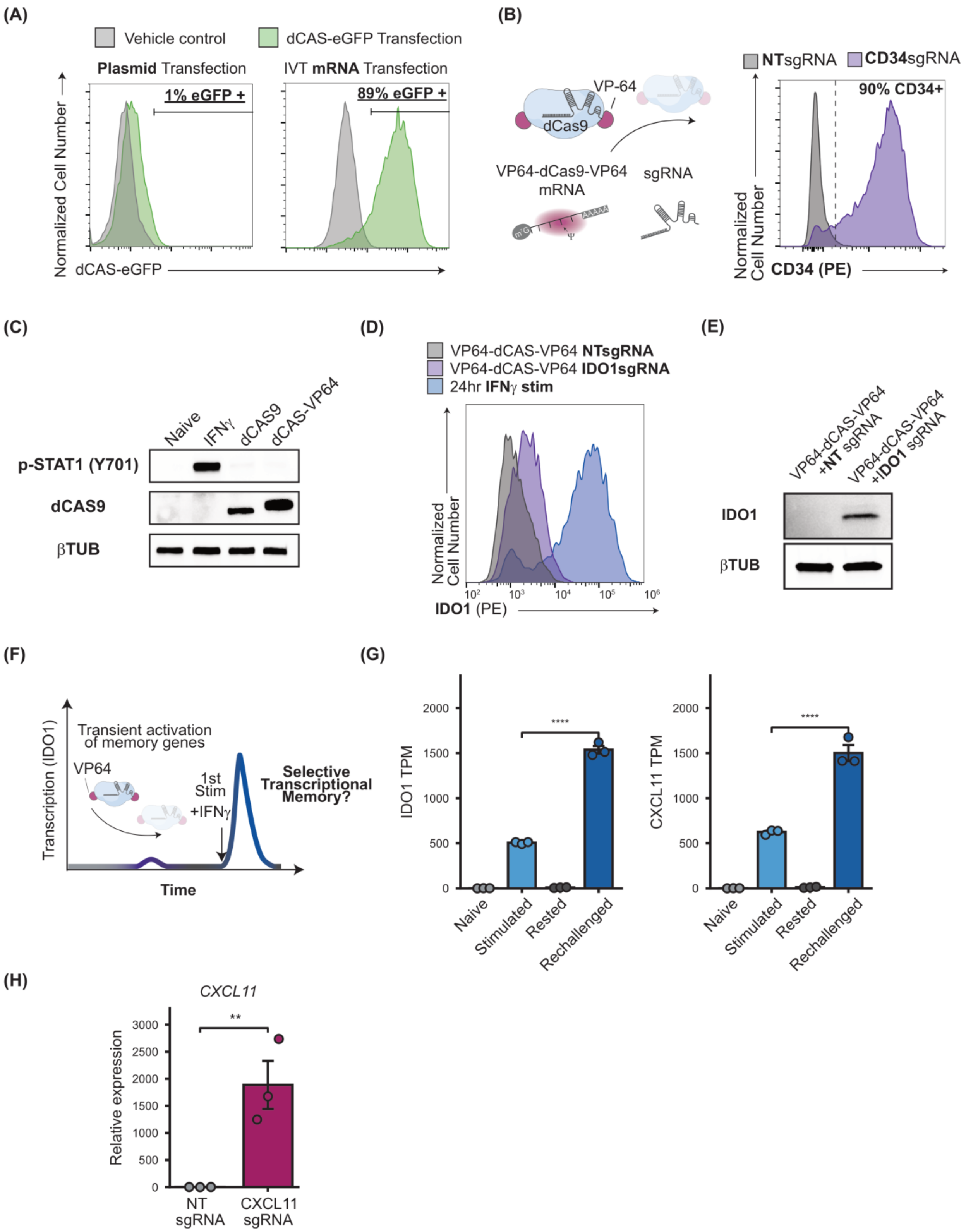

(A)
Vehicle control
dCAS-eGFP Transfection
Plasmid Transfection
IVT mRNA Transfection
1% eGFP +
89% eGFP +
Normalized Cell Number
dCAS-eGFP
(B)
VP-64
dCas9
VP64-dCas9-VP64 mRNA
sgRNA
NTsgRNA
CD34sgRNA
90% CD34+
Normalized Cell Number
CD34 (PE)
(C)
Naive
IFNγ
dCAS9
dCAS-VP64
p-STAT1 (Y701)
dCAS9
βTUB
(D)
VP64-dCAS-VP64 NTsgRNA
VP64-dCAS-VP64 IDO1sgRNA
24hr IFNγ stim
Normalized Cell Number
IDO1 (PE)
(E)
VP64-dCAS-VP64 +NT sgRNA
VP64-dCAS-VP64 +IDO1 sgRNA
IDO1
βTUB
(F)
Transient activation of memory genes
VP64
1st Stim +IFNγ
Selective Transcriptional Memory?
Transcription (IDO1)
Time
(G)
IDO1 TPM
CXCL11 TPM
****
Naive
Stimulated
Rested
Rechallenged
(H)
CXCL11
Relative expression
**
NT sgRNA
CXCL11 sgRNA

**Fig. S5. mRNA-delivered CRISPRa drives efficient, locus-specific activation of *IDO1* and *CXCL11* in MSCs without canonical IFNγ signaling.**
**(A)** Flow cytometry of transfection efficiency comparing plasmid DNA versus in vitro transcribed (IVT) mRNA delivery of a dCas9–eGFP fusion in MSCs; vehicle control (grey) and dCas9–eGFP (green). Left, plasmid; right, IVT mRNA (gate indicated). **(B)** Left, schematic of the CRISPRa system: co-delivery of VP64–dCas9–VP64 mRNA with a synthetic sgRNA. Right, flow cytometry of surface CD34 after VP64–dCas9–VP64 mRNA transfection with a non-targeting (NTsgRNA, grey) or *CD34*-targeting (CD34sgRNA, purple) sgRNA; dashed line, positive gate. **(C)** Western blot on lysates from Naive, IFNγ-stimulated, dCas9-transfected, and VP64–dCas9–VP64-transfected cells, probed for phospho-STAT1 (Y701), dCas9, and βTUB. **(D)** Flow cytometry of intracellular IDO1 for VP64–dCas9–VP64 + NTsgRNA (grey), + IDO1sgRNA (purple), and 24 hr IFNγ (blue). **(E)** Western blot for IDO1 and βTUB (loading) comparing VP64–dCas9–VP64 + NT versus + *IDO1* sgRNA. **(F)** Schematic of the experimental rationale: transient VP64-based CRISPRa activation of a memory gene followed by a first IFNγ stimulation, testing whether transient locus-specific activation installs selective transcriptional memory. **(G)** RNA-seq expression of *IDO1* (left) and *CXCL11* (right) in TPM across the IFNγ training regime (Naive, Stimulated, Rested, Rechallenged). **(H)** RT-qPCR of *CXCL11* relative expression after VP64–dCas9–VP64 + NTsgRNA versus + CXCL11sgRNA. Bars (G, H) show mean ± SEM with individual replicates n = 3. Significance: G, gene-level DESeq2 adjusted P (Rechallenged versus Stimulated); H, Welch's unpaired t-test on ΔCt. **** $P < 0.0001$, *** $P < 0.001$, ** $P < 0.01$, * $P < 0.05$; ns, not significant.

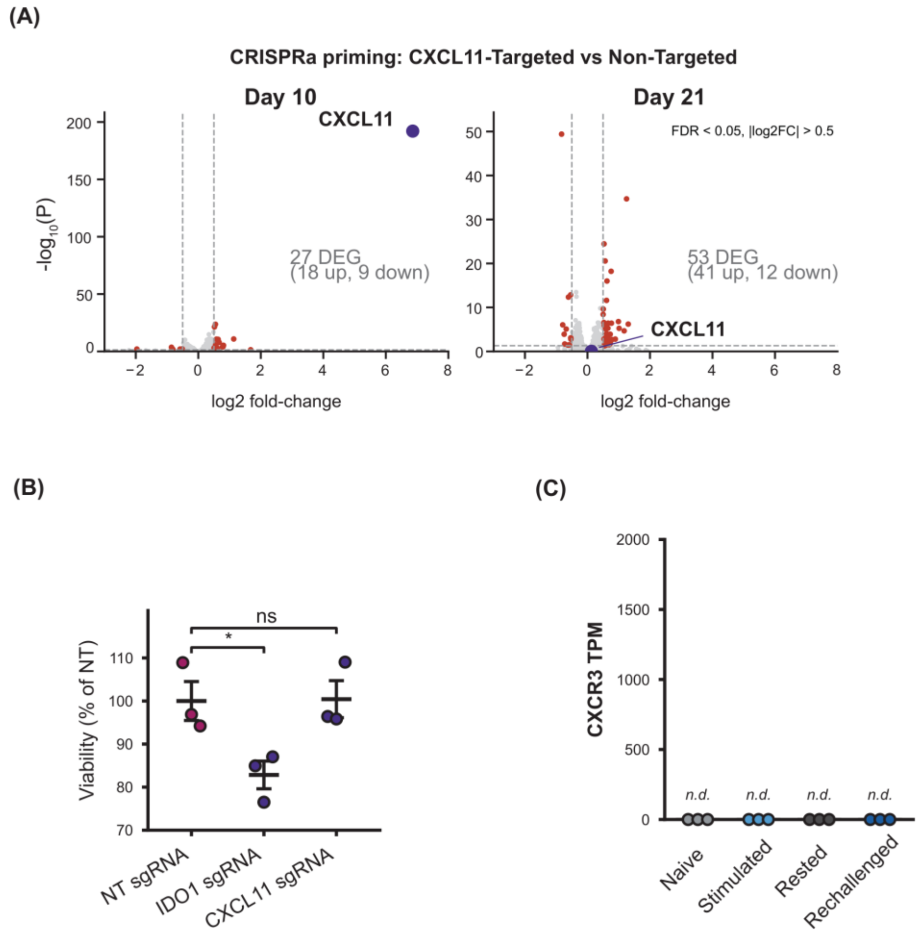


**Fig. S6. *CXCL11*-targeted CRISPRa is minimally perturbative.**
**(A)** Volcano plots of differential expression in the rested state after CRISPRa priming (*CXCL11*-targeted versus non-targeted; −log10 FDR versus log2 fold change) at Day 10 (left) and Day 21 (right); y-axis scale differs between sub-plots. *CXCL11* highlighted in blue; red points, significant DEGs; dashed lines, FDR < 0.05 and |log2FC| > 0.5. **(B)** Alamar Blue cell viability 72 hr after VP64–dCas9–VP64 transfection with NTsgRNA, IDO1sgRNA, or CXCL11sgRNA, normalized to NTsgRNA (set to 100%). **(C)** *CXCR3* transcript abundance (TPM, RNA-seq) across the four IFNγ training conditions; individual replicates shown. *CXCR3*, the receptor for CXCL11, is not detected (n.d.) in any condition. In B, points are individual replicates with mean ± SEM (n = 3 biological replicates); significance by one-way ANOVA with Dunnett's post-test versus NTsgRNA. In A, significance is the gene-level DESeq2 adjusted P (Wald test, Benjamini–Hochberg FDR). **** P < 0.0001, *** P < 0.001, ** P < 0.01, * P < 0.05; ns, not significant.

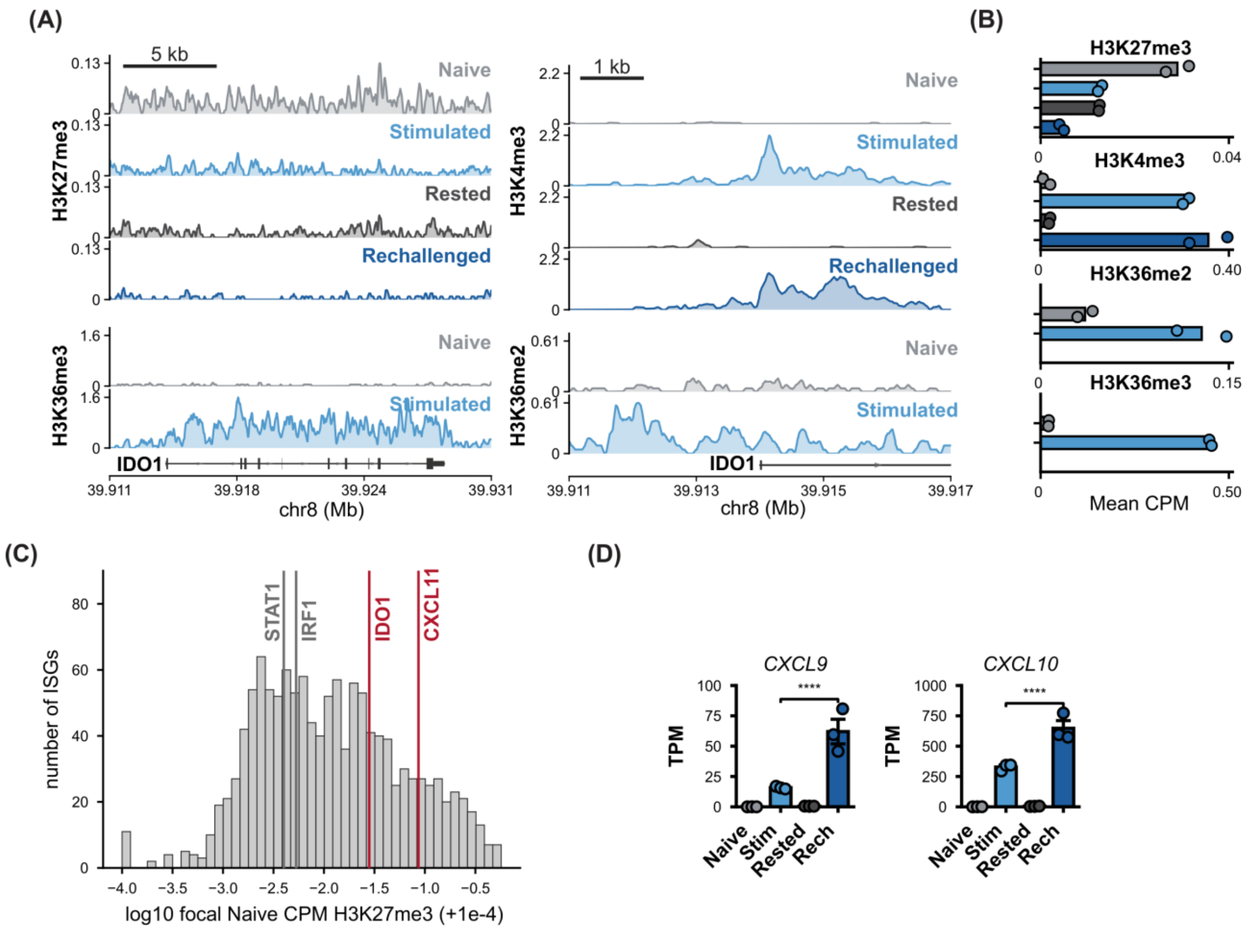

**Fig. S7. H3K27me3 dynamics instruct transcriptional memory at *IDO1* and *CXCL11*.** **(A)** Genome-browser CUT&RUN tracks at the *IDO1* locus across the IFNγ training regime. Left, H3K27me3 and H3K36me3 spanning the gene body. Right, H3K4me3 and H3K36me2 centered on the *IDO1* promoter/TSS. H3K4me3 and H3K27me3 for all four conditions; H3K36me2 and H3K36me3 for Naive and Stimulated only. **(B)** Mean CUT&RUN signal (CPM) at the shown *IDO1* promoter/gene for each mark; n = 2. **(C)** Histogram of promoter H3K27me3 across mappable IFNγ-stimulated (ISG) promoters in the Naive condition. X-axis, log10 focal Naive CPM; vertical lines mark the non-memory genes *STAT1* and *IRF1* (grey) and the memory genes *IDO1* and *CXCL11* (red). **(D)** RNA-seq expression (TPM) of *CXCL9* and *CXCL10*, the co-regulated memory genes adjacent to *CXCL11*, across the four training conditions. Bars, mean ± SEM with individual replicates (n = 3 biological replicates); in D, significance by one-way ANOVA with Tukey's post-test. **** $P < 0.0001$, *** $P < 0.001$, ** $P < 0.01$, * $P < 0.05$; ns, not significant.

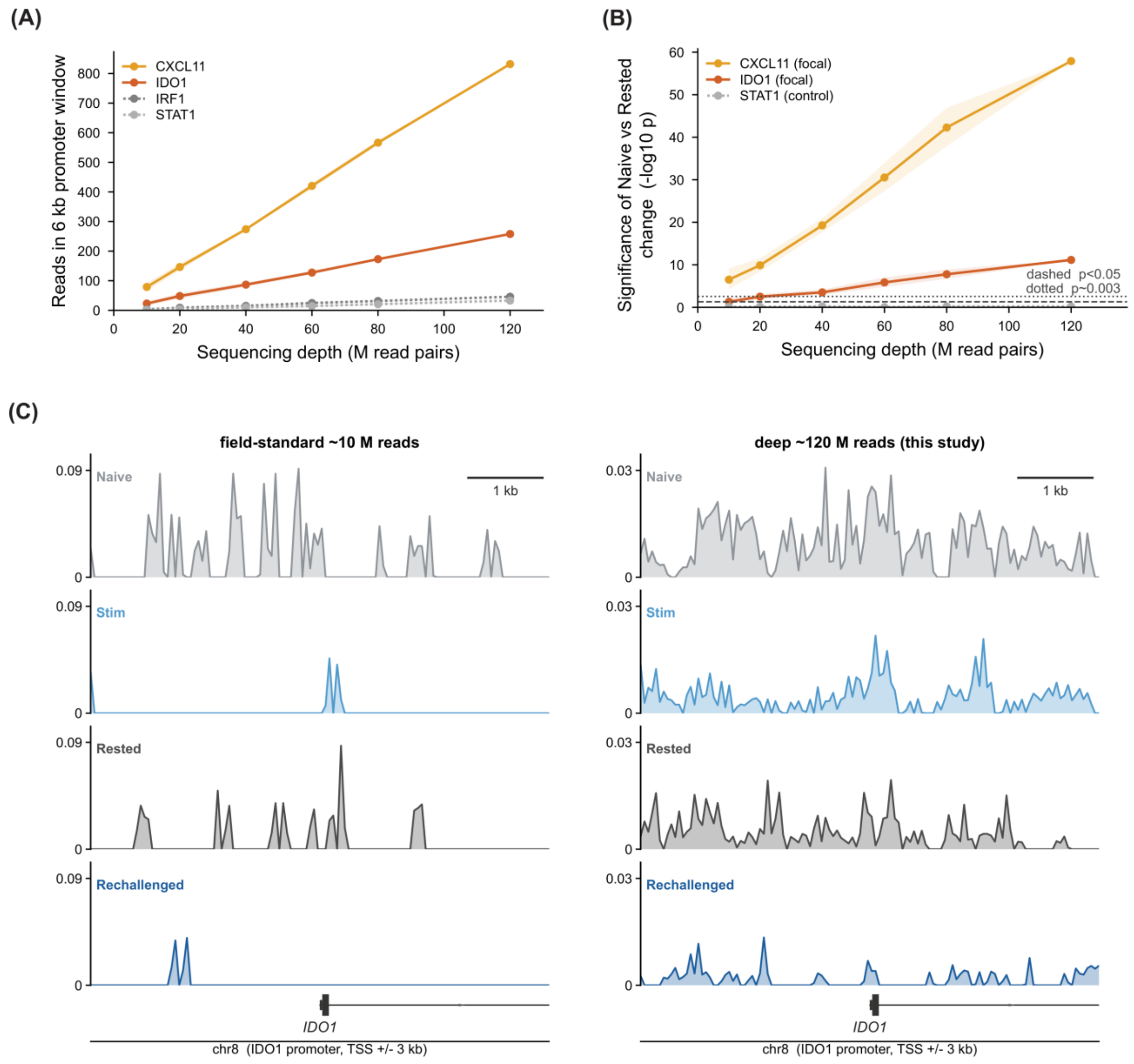


**Fig. S8. Deep sequencing is required to resolve focal H3K27me3 changes at memory-gene promoters.**

**(A)** Read accumulation within a 6 kb promoter window versus H3K27me3 CUT&RUN sequencing depth (M read pairs), for the focal memory loci *CXCL11* (yellow) and *IDO1* (orange) and the non-memory controls *IRF1* (dark grey, dotted) and *STAT1* (light grey, dotted). **(B)** Statistical power to detect the focal Naïve-versus-Rested H3K27me3 change versus sequencing depth (−log10 p), for *CXCL11* (yellow), *IDO1* (orange), and *STAT1* (grey, dotted); shaded bands, variability. Horizontal reference lines, $p < 0.05$ (dashed) and $p \sim 0.003$ (dotted). **(C)** Representative H3K27me3 CUT&RUN tracks across the *IDO1* promoter (chr8, TSS ± 3 kb), comparing field-standard ~10 M reads (left) with the deep ~120 M reads used here (right), for the four training conditions (Naïve, Stim, Rested, Rechallenged).

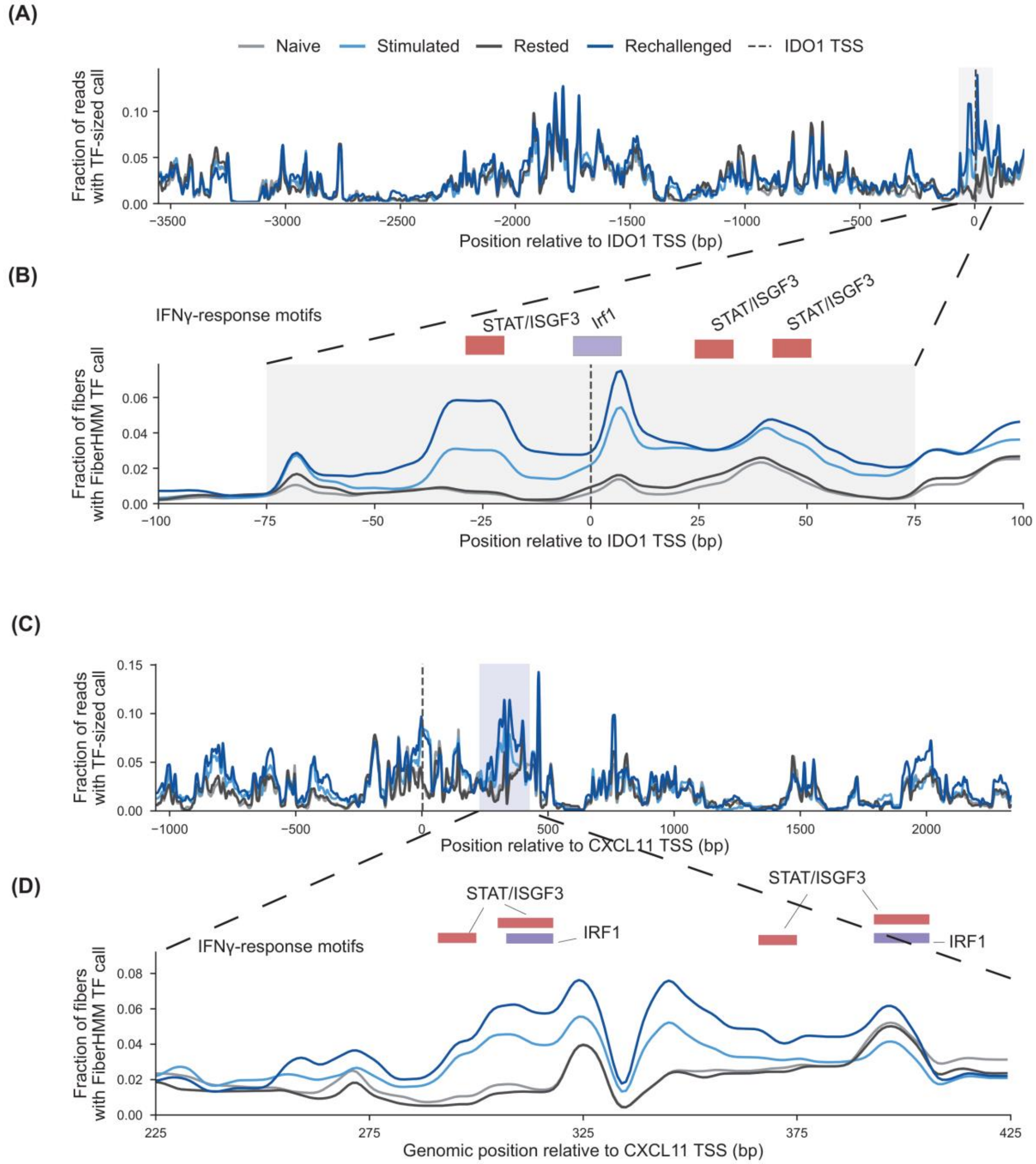


**Fig. S9. Transcription-factor footprint density at the *IDO1* and *CXCL11* promoters is higher on rechallenge than on primary stimulation.**

**(A)** Aggregate density of TF-sized protected footprints called by the FiberHMM model across the *IDO1* amplicon. Shaded band, the promoter-proximal window expanded in B. **(B)** Zoom of the promoter-proximal window (±100 bp of the *IDO1* TSS); same conditions and colors as A. Colored bars denote IFNγ-response motif positions flanking the TSS: STAT/ISGF3 (red) and IRF1 (purple). **(C)** The same aggregate TF-footprint density across the *CXCL11* amplicon, plotted versus position relative to the *CXCL11* TSS; shaded band, the promoter-proximal window expanded in D. **(D)** Zoom of the *CXCL11* promoter-proximal window (~+225 to +425 bp from the TSS); same conditions and colors as C, with STAT/ISGF3 (red) and IRF1 (purple) motif positions marked.

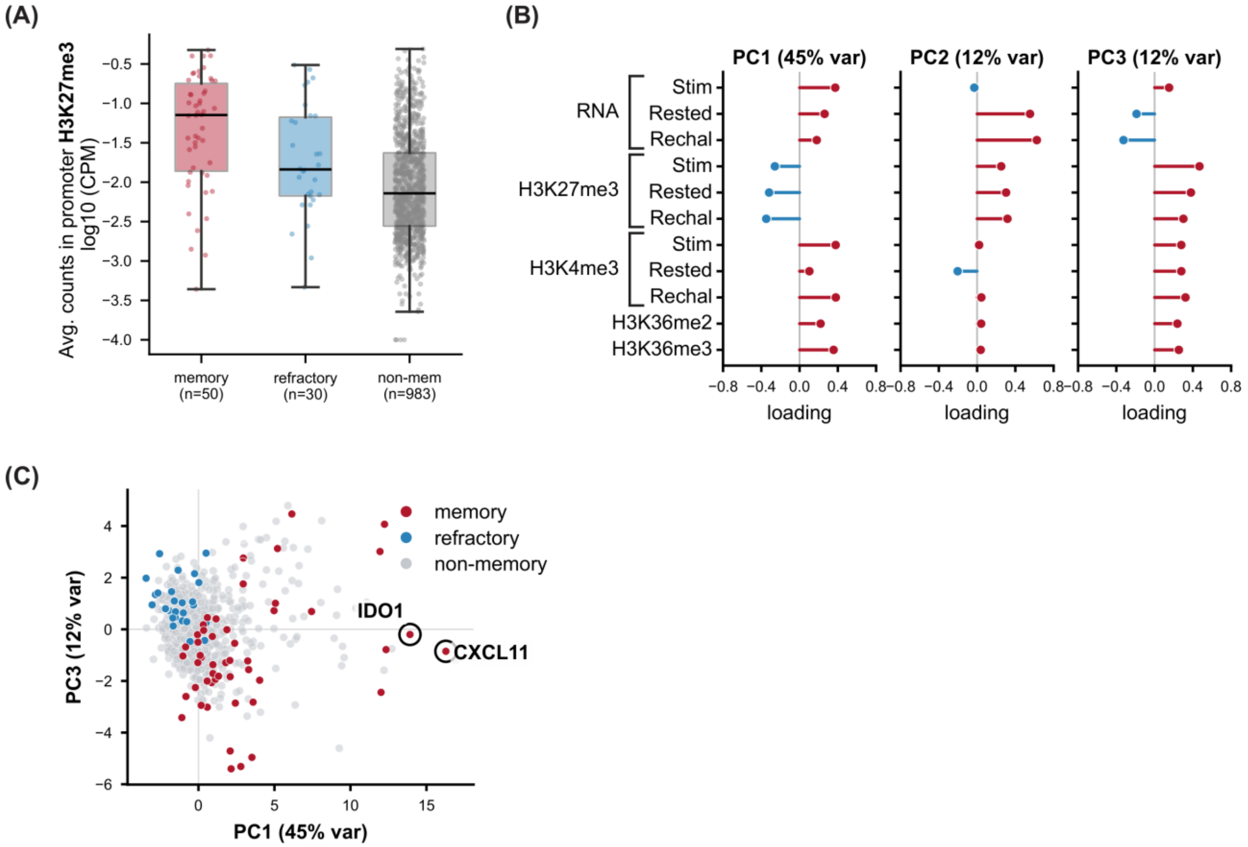


**Fig. S10. Multi-omic PCA Separates highly responsive memory genes.**
**(A)** Boxplots of naive-state promoter H3K27me3 for memory (n = 50), refractory (n = 30), and non-memory (n = 983) genes (log10 average CPM across the promoter window); individual genes overlaid as points. **(B)** Feature loadings from the multi-omic PCA of IFNγ-responsive genes for PC1 (45% of variance), PC2 (12%), and PC3 (12%). Lollipops show the 11 standardized, naive-relative features (RNA at Stim/Rested/Rechal; promoter H3K27me3 and H3K4me3 at Stim/Rested/Rechal; promoter H3K36me2 at Stim; gene-body H3K36me3 at Stim); positive loadings red, negative blue. **(C)** Multi-omic PCA of IFNγ-responsive genes (PC3 versus PC1), colored by classification as in A; *IDO1* and *CXCL11* circled.

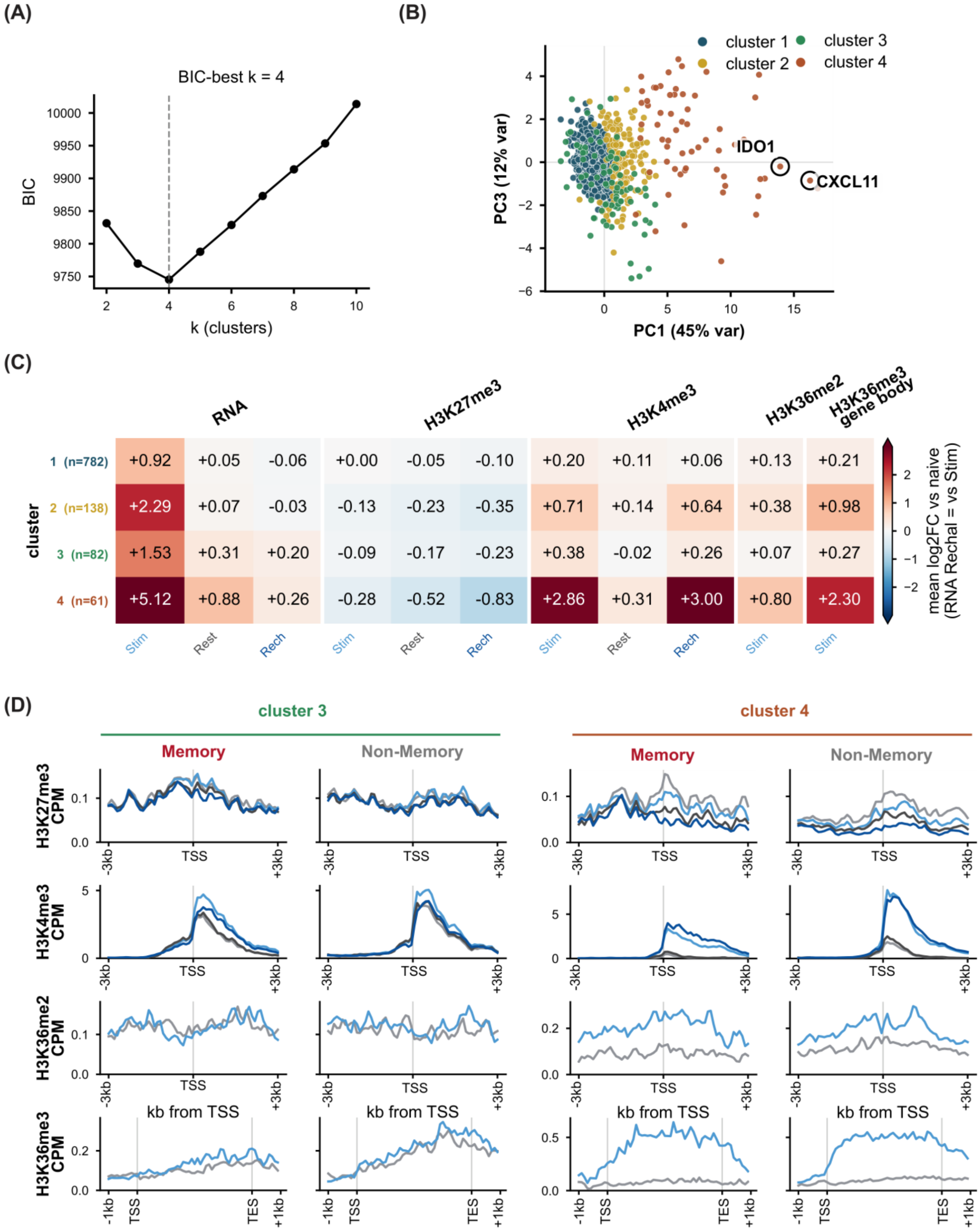
(A)
BIC-best k = 4
BIC
k (clusters)
(B)
cluster 1
cluster 2
cluster 3
cluster 4
IDO1
CXCL11
PC3 (12% var)
PC1 (45% var)
(C)
RNA
H3K27me3
H3K4me3
H3K36me2
H3K36me3 gene body
cluster
1 (n=782) +0.92 +0.05 -0.06 +0.00 -0.05 -0.10 +0.20 +0.11 +0.06 +0.13 +0.21
2 (n=138) +2.29 +0.07 -0.03 -0.13 -0.23 -0.35 +0.71 +0.14 +0.64 +0.38 +0.98
3 (n=82) +1.53 +0.31 +0.20 -0.09 -0.17 -0.23 +0.38 -0.02 +0.26 +0.07 +0.27
4 (n=61) +5.12 +0.88 +0.26 -0.28 -0.52 -0.83 +2.86 +0.31 +3.00 +0.80 +2.30
Stim Rest Rech Stim Rest Rech Stim Rest Rech Stim Stim
mean log2FC vs naive (RNA Rechal = vs Stim)
(D)
cluster 3
cluster 4
Memory
Non-Memory
H3K27me3 CPM
H3K4me3 CPM
H3K36me2 CPM
H3K36me3 CPM
-3kb
TSS
+3kb
kb from TSS
-1kb
TES
+1kb

**Fig. S11. Gaussian-mixture clustering of IFNγ-responsive genes demonstrates H3K27me3 hysteresis delineates highly-responsive memory genes, however non-memory genes can share the chromatin signature.**

**(A)** Gaussian-mixture-model selection: Bayesian information criterion versus number of clusters k (2–10); BIC is minimized at k = 4, the chosen cluster number. **(B)** PC3 versus PC1 colored by GMM cluster assignment (cluster 1, teal; cluster 2, yellow; cluster 3, green; cluster 4, orange); *IDO1* and *CXCL11* circled at high PC1. **(C)** Heatmap of per-cluster mean log2 fold change relative to naive for each feature (rows, clusters 1–4 with n = 782, 138, 82, and 61 genes; columns grouped left to right as RNA, promoter H3K27me3, promoter H3K4me3, promoter H3K36me2, and gene-body H3K36me3); the RNA Rechal column is computed versus Stimulated. Color scale, log2 fold −2 to +2. **(D)** Per-feature metaplots for cluster-3 and cluster-4 genes, each split into memory and non-memory. Rows, top to bottom: H3K27me3 and H3K4me3 across ±3 kb around the TSS (all four conditions), then promoter H3K36me2 (±3 kb) and gene-body H3K36me3 (scaled TSS–TES with ±1 kb flanks) for Naïve and Stimulated only. Cluster 3, n = 32 memory and 31 non-memory; cluster 4, n = 13 memory and 48 non-memory.

**Table S1. Oligonucleotide primers used for quantitative PCR.**
Primers used for reverse-transcription qPCR (RT-qPCR) of transcript abundance, chromatin immunoprecipitation qPCR (ChIP-qPCR) of VP64-dCas9-VP64 occupancy, and CUT&RUN-qPCR of H3K27me3 enrichment. All primers target the human genome or transcriptome and were used with Luna Universal qPCR Master Mix (SYBR detection) on a Bio-Rad CFX real-time system. RT-qPCR expression was normalized to *ACTB* and quantified by the 2^(−ΔΔCt) method; ChIP-qPCR enrichment is reported as percent input; CUT&RUN-qPCR enrichment is reported as fold change relative to the HOX control locus. Internal oligo identifiers (oGB…) are given where available. Genomic coordinates refer to hg38.

| Assay / target | Oligo ID | Forward primer (5'→3') | Reverse primer (5'→3') | Amplicon / template | Figure(s) |
|---|---|---|---|---|---|
| ***RT-qPCR (transcript abundance)*** | | | | | |
| *IDO1* | — | ACGGGACACTTTGCTAAAGGC | GGTTGCCTTTCCAGCCAGACA | cDNA | Fig. S1C; Fig. 3 (EZH2i) |
| *CXCL11* | oGB111 / oGB112 | AAGGACAACGATGCCTAAATCCC | CAGATGCCCTTTTCCAGGACTTC | cDNA | Fig. S5H |
| *IRF1* | — | CTCTCCCCGACTGGCACATC | CCGACTGCTCCAAGAGCTTCA | cDNA | Fig. 3F |
| *STAT1* | — | ATGGCAGTCTGGCGGCTGAATT | CCAAACCAGGCTGGCACAATTG | cDNA | Fig. 3F |
| *ACTB* (reference gene) | — | CACCATTGGCAATGAGCGGTTC | AGGTCTTTGCGGATGTCCACGT | cDNA | All RT-qPCR panels |
| ***ChIP-qPCR (VP64-dCas9-VP64 occupancy)*** | | | | | |
| *IDO1* promoter, sgRNA target site | oGB252 / oGB253 | AATTTCTCACTGCCCCTGTG | CTTATAATAGTTGCTGCCACAGCC | chr8 (*IDO1* promoter); 60 bp | Fig. 2E |
| ***CUT&RUN-qPCR (H3K27me3 enrichment)*** | | | | | |
| *IDO1* promoter (CR_4) | oGB256 / oGB257 | TGCCCCTGTGATAAACTGTGG | CCCCTCAGTGTCTGAAGAGTT | chr8:39,913,819–39,913,899 (81 bp) | Fig. 3 |
| *CXCL11* promoter (CR_1) | oGB260 / oGB261 | ACTGTCCCTTCCTATTTGCAGT | GGCTGGTTACCATCTGGGTT | chr4 (*CXCL11* promoter); 79 bp | Fig. 3 |
| *STAT1* promoter (CR_1) | oGB264 / oGB265 | GGTATTTCCGCCGGCTTCC | AAACAGAACGCCAGTTCCCA | chr2 (*STAT1* promoter); 90 bp | Fig. 3 |
| *IRF1* promoter (CR_1) | oGB268 / oGB269 | ACACTTAGCGGGATTCCCCA | GAAATGACGGCACGCAGC | chr5 (*IRF1* promoter); 87 bp | Fig. 3 |
| HOX control locus (CR_1) | oGB232 / oGB233 | CCAATACTCAGGCGCAGAGA | GTGGGAGACTCAGTTCAGGC | HOX cluster; normalization control | Fig. 3 |

**Table S2. Amplicon primers used for DAF-seq.**

Primers used to generate the targeted promoter amplicons for deaminase-assisted single-molecule fiber sequencing (DAF-seq) with repliQa HiFi ToughMix. Primers carry degenerate G-to-R (G or A) substitutions, introduced during design to reduce G-runs and improve the deamination readout; sequences are given as synthesized (IDT). Coordinates refer to hg38.

| Assay / target | Oligo ID | Forward primer (5'→3') | Reverse primer (5'→3') | Amplicon / template | Figure(s) |
|---|---|---|---|---|---|
| ***DAF-seq amplicon primers*** | | | | | |
| *IDO1* | oGB309 / oGB285 | TCCCATCTACAAARCCATCTCC | TCTCACCACTCTCCRTATCCA | chr8:39,910,187–39,914,251 (4,064 bp) | Fig. 4; Fig. S8 |
| *CXCL11* | oDY002 / oDY007 | TCCCCCARARACATCTAAAAACA | RCTCCAARAACACATTCCTTTTACT | chr4:76,034,865–76,038,554 (3,706 bp) | Fig. 4; Fig. S8 |
| *STAT1* | oGB294 / oGB303 | TCAACCACACCTCTRCATACC | ATAACTCCCACAATTCCCACCT | chr2:191,013,310–191,016,238 (2,928 bp) | Fig. 4 |

**Table S3. Materials and resources.**

Reagents, antibodies, kits, instruments and software used in this study, grouped by category and listed in the order the Materials and Methods introduce them. Catalog numbers are those given by the supplier at the time of purchase. Antibody dilutions and reagent working concentrations are given where the Materials and Methods specify them. Oligonucleotide sequences are given in tables S1 and S2. Entries marked “not stated” are reagents used in this study for which a catalog number or supplier is not specified in the Materials and Methods.

| Reagent or resource | Source | Identifier | Used for |
|---|---|---|---|
| **Biological samples** | | | |
| Human adipose-derived mesenchymal stromal cells, normal | ATCC | PCS-500-011 | All experiments; donor lots 80921243, 80705221, 70040251; used at passage 4 to 7 |
| **Antibodies** | | | |
| IDO1 (clone V1NC3IDO) | eBioscience/Invitrogen | 14-9750-82 | Western blot |
| IDO1, FITC conjugate (clone eyedio) | eBioscience | 11-9477-42 | Flow cytometry |
| IDO1, Alexa Fluor Plus 647 conjugate | eBioscience | 740095MP647 | Flow cytometry |
| CD34, Alexa Fluor 647 | BioLegend (via Fisher) | 50-207-6593 | Flow cytometry |
| Phospho-STAT1 (Y701) | Invitrogen | MA5-15071 | Western blot |
| STAT1, total | Proteintech | 66545-1-Ig | Western blot |
| IRF1 (clone BLR039F) | Abcam | ab243895 | Western blot |
| Cas9/dCas9 | Diagenode | NC1554064 | Western blot |
| Cas9 (clone 4G10) | Diagenode | C15200216-100 | ChIP-qPCR, 5 µg per IP |
| CXCL11 | R&D Systems | MAB672SP | Western blot |
| H3K27me3 | Millipore | ABE44 | Western blot |
| Histone H3, total | Abcam | ab1791 | Western blot loading control |
| Normal IgG isotype control | Invitrogen | A10630 | ChIP-qPCR, 5 µg per IP |
| Anti-rabbit IgG, HRP-conjugated | Cell Signaling Technology | 7074 | Western blot, 1:2,000 |
| Anti-mouse IgG, HRP-conjugated | Cell Signaling Technology | 7076 | Western blot, 1:2,000 |
| hFAB Rhodamine anti-tubulin | Bio-Rad | 12004165 | Western blot loading control |
| H3K4me3 | EpiCypher | 13-0060 | CUT&RUN |
| H3K27me3 | EpiCypher | 13-0055 | CUT&RUN; CUT&RUN-qPCR |
| H3K36me2 | EpiCypher | RD130056 | CUT&RUN |
| H3K36me3 | EpiCypher | 13-0058 | CUT&RUN |
| **Chemicals, peptides and recombinant proteins** | | | |
| DMEM, low glucose (1 g/l) with L-glutamine and sodium pyruvate | Corning | MT10014CV | Culture medium base |
| Fetal bovine serum | MilliporeSigma | F2442 | 5% in culture medium |
| Recombinant human FGF2 | R&D Systems | CFBTFGFB020 | 5 ng/ml in culture medium |
| Recombinant human EGF | R&D Systems | 236-EG-200 | 5 ng/ml in culture medium |
| Penicillin/streptomycin | Gibco | 15-140-163 | 1% in culture medium |
| Recombinant human IFNγ | STEMCELL Technologies | 50-197-6285 | 50 ng/ml, IFNγ training paradigm |
| Recombinant human TNFα, HEK293-expressed, carrier-free | R&D Systems | 10291-TA | 10 ng/ml, TNFα training paradigm |

| Reagent or resource | Source | Identifier | Used for |
|---|---|---|---|
| Tazemetostat (EPZ-6438), EZH2 inhibitor | APExBIO | A8221 | 1 µM for 6 days, with 0.1% DMSO vehicle control |
| Opti-MEM I reduced-serum medium | Gibco | 31985070 | Transfection diluent |
| HindIII-HF | New England Biolabs | R3104S | IVT template linearization |
| RNA Loading Dye (2X) | New England Biolabs | B0363S | IVT product QC gel |
| ssRNA ladder | New England Biolabs | N0362S | IVT product QC gel |
| RIPA lysis and extraction buffer | Thermo | 89900 | Whole-cell lysis |
| Halt Protease Inhibitor Cocktail (100×) | Thermo | 78430 | Whole-cell lysis |
| Mini-PROTEAN TGX precast gels, 7.5% | Bio-Rad | 4561023 | Western blot |
| EveryBlot Blocking Buffer | Bio-Rad | 12010020 | Western blot blocking |
| Clarity Western ECL substrate | Bio-Rad | 1705061 | Western blot detection |
| SuperSignal West Femto substrate | Thermo | 34094 | Western blot detection |
| Protein G Dynabeads | Thermo Fisher | 10003D | ChIP-qPCR capture |
| GolgiPlug (brefeldin A) | BD Biosciences | 555029 | Intracellular CXCL11 accumulation |
| SsDddA single-stranded DNA cytidine deaminase | Stergachis laboratory | not commercial | DAF-seq; 2 µl of 100 µM, 25 °C for 10 min |
| **Critical commercial assays and kits** | | | |
| Cytofix/Cytoperm | BD Biosciences | 554714 | Flow cytometry fixation and permeabilization |
| BCA Protein Assay Kit | Thermo | 22327 | Protein quantification |
| Trans-Blot Turbo RTA Mini LF PVDF Transfer Kit | Bio-Rad | 1704274 | Western blot transfer |
| RNeasy Plus Mini Kit | Qiagen | 74136 | RNA extraction |
| iScript Advanced cDNA Synthesis Kit | Bio-Rad | 1725038BUN | cDNA synthesis, 250 ng RNA |
| Luna Universal qPCR Master Mix | New England Biolabs | M3003E; M3003 | RT-qPCR, ChIP-qPCR, CUT&RUN-qPCR |
| Cloning Kit for mRNA Template | Takara | 6143 | T7 IVT template construction |
| CleanCap AG (3' OMe) CleanScript IVT Kit | TriLink BioTechnologies | K-7413 | In vitro transcription of CRISPRa mRNA |
| CRISPRevolution sgRNA EZ Kit, 1.5 nmol | Synthego | not stated | Chemically modified sgRNA synthesis |
| Lipofectamine MessengerMAX | Invitrogen | LMRNA015 | mRNA and sgRNA reverse transfection, 1.5 µl/ml |
| MinElute PCR Purification Kit | Qiagen | 28004 | ChIP-qPCR DNA cleanup |
| CUTANA ChIC/CUT&RUN Kit | EpiCypher | 14-1048 | CUT&RUN |
| Nuclei Extraction Buffer | EpiCypher | 21-1026 | CUT&RUN and DAF-seq nuclei isolation |
| CUTANA CUT&RUN Library Prep Kit | EpiCypher | 14-1002 | CUT&RUN library preparation |
| DNeasy Blood & Tissue Kit | Qiagen | 69506 | DAF-seq genomic DNA extraction |
| repliQa HiFi ToughMix | Quanta Bio | 95200 | DAF-seq promoter amplicon PCR |
| NEBNext Ultra II RNA Library Prep Kit for Illumina | New England Biolabs, via Azenta/GENEWIZ | not stated | RNA-seq library preparation, poly-A selection |
| **Oligonucleotides** | | | |
| *IDO1* sgRNA "sg5", protospacer 5'-GTGACCACAGTTTATCACAG-3' | Synthego | not stated | CRISPRa targeting of *IDO1* |

| Reagent or resource | Source | Identifier | Used for |
|---|---|---|---|
| *CXCL11* sgRNA, protospacer 5'-AAGCACTGTGAAAACCCAGA-3' | Synthego | not stated | CRISPRa targeting of *CXCL11* |
| *CD34* sgRNA, protospacer 5'-GAAAGCTGAACGAGGCATC-3' | Synthego | not stated | CRISPRa benchmark, chr1:207,911,360-207,911,378 |
| Negative control (scramble) sgRNA #1 | Synthego | 063-1010-000-000 | Non-targeting CRISPRa control |
| qPCR primers | Integrated DNA Technologies | table S1 | RT-qPCR, ChIP-qPCR and CUT&RUN-qPCR |
| DAF-seq amplicon primers, degenerate G-to-R | Integrated DNA Technologies | table S2 | DAF-seq promoter amplicons |
| **Instruments and sequencing** | | | |
| CFX96 Touch Deep Well real-time system | Bio-Rad | 3600037 | All qPCR (CFX Maestro v5.2.008.0222) |
| NovaSeq X Plus | Illumina, via Admera Health | not stated | CUT&RUN sequencing, 2 × 150 bp |
| Oxford Nanopore sequencing | Oxford Nanopore | not stated | DAF-seq, approximately 5,000 reads per sample |
| **Software and algorithms** | | | |
| STAR | Dobin et al. | v2.7.11b | RNA-seq genome index and alignment |
| RSEM | Li and Dewey | v1.3.3 | RNA-seq alignment and quantification |
| tximport | Bioconductor | v1.36.0 | Transcript-level import |
| DESeq2 | Bioconductor | v1.50.2 | Differential expression |
| clusterProfiler; enrichplot | Bioconductor | v4.16.0; v1.28.0 | Gene-set enrichment |
| fastp | Chen et al. | v0.24.0 | CUT&RUN read trimming |
| Bowtie2 | Langmead and Salzberg | v2.5.4 | CUT&RUN alignment to hg38 |
| bedtools | Quinlan and Hall | v2.31.1 | Interval operations |
| deepTools | Ramírez et al. | v3.5.5 | Coverage tracks and metaplots |
| minimap2 | Li | v2.28 | DAF-seq amplicon alignment |
| fibertools-rs | Stergachis laboratory | v0.7.0 | DAF-seq deamination decoration |
| FiberHMM | Stergachis laboratory | v2.10.5 | MSP and TF footprint calling |
| R | R Core Team | v4.5.1 | Statistical analysis |
| Python | Python Software Foundation | v3.13; v3.12 | Analysis and figures (matplotlib v3.10.8, numpy v2.2.3, scipy v1.15.2, pysam v0.23.0, statsmodels) |
